# Musicality in protein interaction dynamics informs the multi-scale evolution of prosocial behavior

**DOI:** 10.1101/2025.03.13.643054

**Authors:** Gregory A. Babbitt, Lin Wang, Ernest P. Fokoue

## Abstract

Like animal vocalization and display, human singing and dancing allows non-verbal establishment of behavioral co-relation (i.e. correlation) between individuals. The predictable mathematical structure of music is its most defining acoustic property, allowing human synchronization of both physical behavior and emotion. In the biomolecular world, some proteins also interact in groups to achieve strong spatiotemporal co-relationships. This is prominent in amyloids, where many disordered fibrils individually conform to overall solenoid structures. We hypothesize that the vibrational frequencies captured during amyloid protein interactions may also exhibit elements of musicality related to this form of prosocial behavior. Here, we develop a non-abstract data sonification method for computer-simulated molecular dynamic interactions. We apply auto-correlational and spectral cross-correlational analyses to a collection of sounds, defining 11 acoustic features that allow accurate machine learning classification of music from other types of natural sounds. By analyzing statistical shifts in these correlative features defining musicality, we demonstrate that amyloid interactions are more speech-like and musical than less structurally conforming protein interactions, primarily due to significant shifts in memory (persistence) and first order autocorrelation. We also find that music has less feature shift away from animal vocalization than human speech, suggesting it may have predated the evolution of language.

## Introduction

Eugene Wigner once famously questioned whether mathematics was a purely human invention, like specific languages, or whether it was a reflection of a universal physical reality (Wigner, 1960). We could ask similar questions about music. Is it purely a product of human culture and creativity, or does it also reflect something more universal that exists in nature beyond the human mind. Perhaps like mathematics, musical sounds probably contain feature elements with both internal and external representation. Internally, musical sounds have a large excitatory effect on the neurophysiology of the human brain (Sacks, 2008; Trimble and Hesdorffer, 2017), and externally, musical sounds have a variable and yet predictable mathematical structure in temporal properties, similar to human speech and other animal vocalizations (Kello et al., 2017). While human speech and music are similar in their acoustics, they also differ in their regularity of amplitude modulation (Chang et al., 2024) and narrowness of pitch range (Wolfe, 2002), with music being more regular and stable than speech. These features of musical acoustics may provide potential clues to its evolutionary function. The ability of music to stimulate the motor cortex through cadence and rhythm (Gordon et al., 2018; Izbicki et al., 2020), combined with stimulation of affective empathy (perhaps via ‘mirror neurons’) in the premotor cortex (Bedoya et al., 2021; Gallese et al., 2011; Kilner and Lemon, 2013; Trost et al., n.d.), suggest an ancestral role of music could have been for the non-verbal organization of group social behavior in time and space (Spitzer, 2021).

But is musicality a purely human adaptation? What evidence is there that musicality exists outside of humans? Recently, biomusicologists (Fitch, 2015, 2006; Honing, 2018; Honing et al., 2015; Merker et al., 2015; Ravignani et al., 2014) have identified features of human musicality in our close primate relatives, including isochrony and improvisational duetting, (De Gregorio et al., 2024; Raimondi et al., 2023). Additionally, the avian syrinx, which is more complex than the mammalian larynx, is also capable of generating sounds that exhibit some degree of musicality (Baptista and Keister, 2005; Bilger et al., 2021; Rothenberg et al., 2014). Biomusicologists have also discovered that a diverse range of animals are behaviorally responsive to music when it is acoustically adjusted for them (Snowdon and Teie, 2009) again suggestive of a common neurophysiological function for music. In most animals, specific patterns of vocalization in combination with specific physical motions (i.e. performance displays), are a common means of conveying fitness, establishing territorial boundaries, attracting potential mates, strengthening pair-bonds, and organizing social groups. In most animals, nonaggressive interactions are rare outside of immediate kinship. Humans are one of only a few species of mammals that have crossed over into highly evolved prosocial behavior, where unrelated individuals cooperate to achieve group goals (Crespi and Yanega, 1995). This would require the evolution of neurophysiological mechanisms that can organize human group activity beyond that of most other mammals. In social encounters with strangers, the predictability of music and dance may have provided a useful means for building social cohesion (Trainor, 2015; Trehub et al., 2015), by limiting the threat of physical aggression while also assessing the fitness of whole groups at the same time. Because the human larynx can produce a much larger range of sound than is necessary for speech, some have also hypothesized that its ancestral evolution was driven by a need to produce a greater variety of non-verbal sounds than are required to convey information through language (*When Bjork Met Attenborough*, 2014). Neural overlap in the processing of music and speech might also suggest such a common ancestral origin (Peretz et al., 2015). Unfortunately, because music was not notated and recorded until modern times, and because archeological evidence of musical instruments offers little societal context, the evolutionary origin of music remains quite speculative (Spitzer, 2021). However, comparative analysis of the acoustic features of music, human speech, animal vocalization, in the context of other natural sounds, could prove useful in identifying what acoustic aspects of musicality are genetically ancestral vs. culturally derived (Brown et al., 2014; Trehub et al., 2015). Recent advances in machine learning can also allow identification of the relative importance of different acoustic features of music without the inherent subjectivity that arises from the human listener.

Complex systems with highly socially interactive components are not restricted to human groups. Many other highly organized physical and biological systems exist at all scales in nature (Jensen, 1998; Sornette, 2006). One might then wonder if the mathematically regular and predictable acoustic structure of music might also manifest itself across various scales within complex systems in general. Could musicality in biological systems exist at smaller scales than that of organisms? In the biomolecular world, proteins share many of the same behaviors of human musical performers and/or dancers. Like all molecules, proteins produce vibrational frequencies, and because of their large size and soft matter composition, they produce a large range of complex dynamics. Proteins are often highly specific in their interactions with each other, much like mate pair bonding in the animal world. They also compete to establish biochemical co-relations that support such interactions. Like organisms, proteins often have bilateral or radial symmetry and produce non-random directed motions similar to a choreographed display (Babbitt et al., 2024). Some proteins even form specific co-relational structures with each other to form larger functional groups. In amyloids, individual fibrils convert from intrinsically disordered states into groups with highly repetitive solenoid configurations, where each individual fibril conforms its structure and dynamics to that required by the larger group (Chatani et al., 2021; Mohd Nor Ihsan et al., 2023). Such behavior in the molecular world is similar to behavior of individuals in highly conforming prosocial/eusocial systems, such as social insects (Wilson and Wilson, 2007; Wilson and Hölldobler, 2005; Wilson and Nowak, 2014). Our analogy across biological scales is outlined in Figure 1. While current definitions of eusociality assume individual costs/benefits to genetic fitness (Crespi and Yanega, 1995; Pfattheicher et al., 2022) obviously absent at the level of proteins, the ‘fitness’ of a protein aggregation might be measured via alternative currencies, such as free energy or thermostability. The existence of eusociality (i.e. true self-sacrifice) amongst social castes in the animal world has been the subject of decades long debate, particularly where humans are concerned (Hooper et al., 2015; Kay et al., 2020). However, the existence of human prosocial behavior is better documented if not completely understood (Melis, 2018; Pfattheicher et al., 2022; Silk and House, 2011). Here we hypothesize that if musicality is a proximate mechanism for prosocial cohesion in the human-animal world, perhaps it might also be evident in highly cohesive biomolecular systems, such as amyloids, as well.

**Figure 1.**
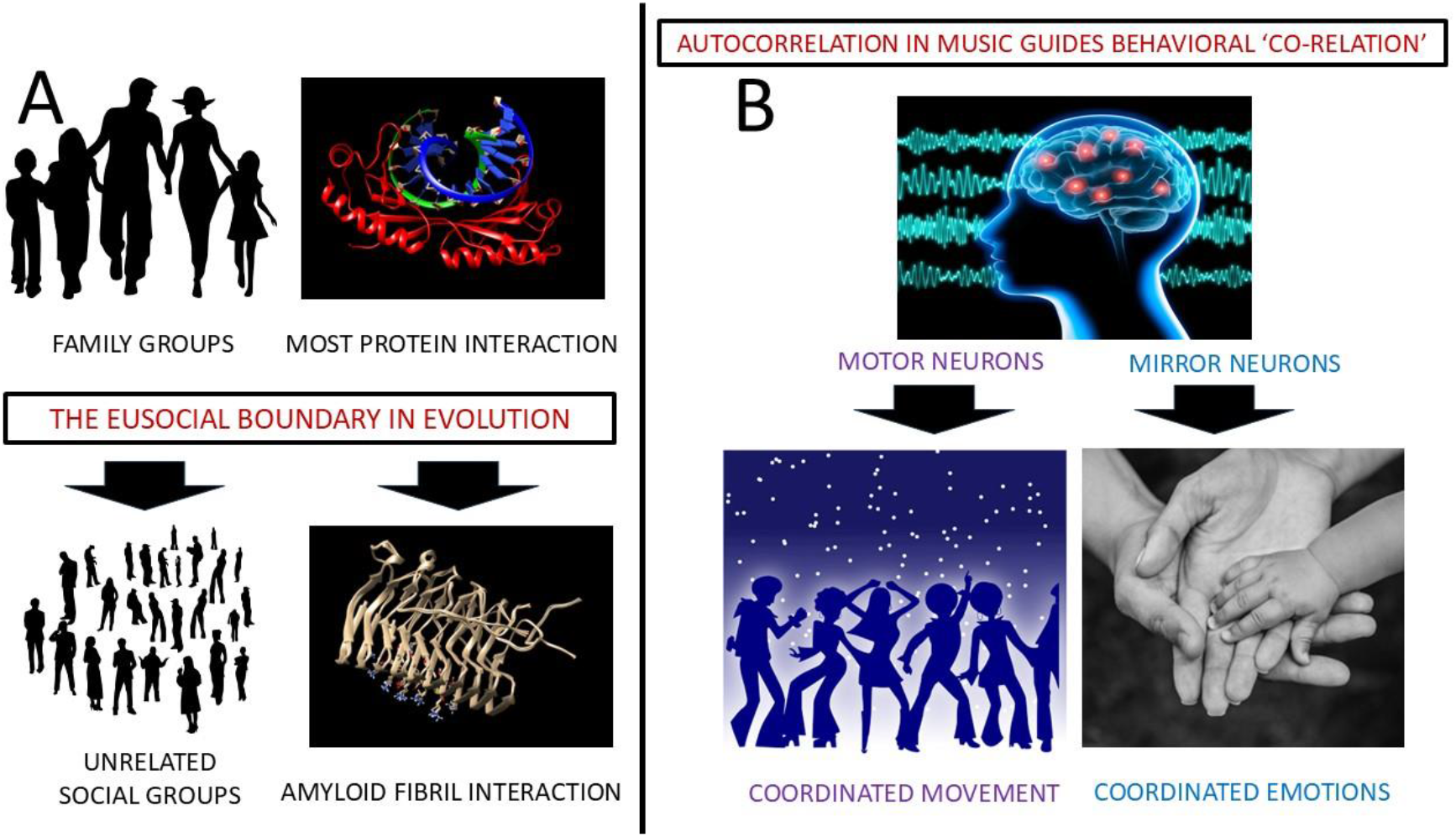
The central hypothesis of molecular biomusicology. conjectures that (A) at an ultimate level, the evolution of prosocial behavior at both the organismal level and molecular level requires high levels of conforming behavior. (B) At a proximate level, prosocial evolution might require autocorrelational cues that can align spatial-temporal activity (as well as emotion in the higher organisms). Thus, some elements or features of music, which is highly autocorrelative, may reside in nature outside of human culture in prosocial or eusocial systems at many scales.

To investigate this idea, we first need to develop a method of data sonification that can be applied in a concrete, non-abstracted way to computer simulated motions of protein interactions (i.e. molecular dynamics simulations). Second, we need some measurable mathematical properties of musicality that can be identified acoustically. A fundamental mathematical measure of co-relationship is the correlation coefficient (i.e. Galton’s literal origin of the term (Galton, 1888)). Sounds can be measured in both the frequency and time domains producing data capturing the self-similar (i.e. autocorrelated) elements of musical structure that determine melodic sequence complexity, harmonic layering (and progression), and cadence/rhythm (or lack thereof). Some previous studies have derived musical abstraction from protein sequences, but to our knowledge, none of these studies have attempted to examine an explicit role for musicality in the context of protein function. Here we developed the Atomdance AudioVisualizer (AAV), a python-based data sonification software pipeline based upon our previously published ATOMDANCE software suite for comparative protein dynamics (Babbitt et al., 2024). The generation of sound files as non-abstract representations of protein interactions via the AAV allows us to further analyze their musicality (or lack thereof) in the context of other natural sounds.

## Methods overview

The Atomdance AudioVisualizer (AAV) is a 25+ step computational pipeline for converting protein interaction dynamics into sound representation. The AAV uses conventional molecular dynamics (MD) simulations to create input representing two states of dynamic comparison; A) a protein/nucleic acid/ligand bound dynamic state, and B) an unbound dynamic state. The AAV also produces a much shorter highly sampled MD run from which vibrational frequencies of individual amino acid are extracted following the method of (Wang, 2019). These frequencies are later adjusted to the pitch sensitivity of the human ear as a function of the amount of atom fluctuation in the long range MD simulations in the bound state. The AAV then employs choreographic analysis to define regions of coordinated site dynamics (Babbitt et al., 2024) which are subsequently utilized for the overlaying of amino acid frequencies according to their intrinsically coordinated group dynamics (i.e. establishing choir sections). Lastly, kernel-based denoising is employed to define site-wise maximum mean discrepancy (MMD) in atom fluctuations when comparing dynamics of proteins in their bound and unbound states (Babbitt et al., 2024). Because most of the dynamics in MD simulation is noise generated by random collisions with the surrounding solvent, and because machine learning cannot learn from noise, the MMD represents the component of motion that is functional (i.e. nonrandom) at a given site. High positive or negative values of MMD effectively identify amino acid sites that are momentarily active when important functions involving protein interactions occur. These can include dampened fluctuations due to site-specific binding (i.e. negative MMD), or amplified fluctuation due to activation of loop regions (positive MMD). The AAV uses this temporal information about site function to determine which amino acid ‘choirs’ are vocally active at any given time (i.e. potentially establishing sequences of note with potentially melodic and/or rhythmic/harmonic characteristics). The intervals between these notes are rendered in two ways. The first rendering uses a fixed interval of 0.25 seconds producing a potentially more musical sounding sequence. The second rendering uses a variable time interval where the length of the note is inversely proportional to the momentary strength of binding during the MD simulation. This is determined by the area under the MMD curve across the length of the protein. This second acoustic rendering of protein interaction produces a potentially more speech-like sequence of sound. An overview of the AAV pipeline for converting protein interaction dynamics into sound is shown graphically in Figure 2A and schematically in Figures 2B-2C. All math formulae are in the equations in the Supplemental Methods section referenced in the schematic overviews. The AAV produces sound and movie files of protein interaction dynamics that can later be analyzed to investigate the roles of different features of musicality in the biomolecular world. Unlike most other previous computational approaches to converting protein information into sound (Takahashi and Miller, 2007; Tay et al., 2021; Yu et al., 2019), the AAV pipeline does not make use arbitrary assignment of sounds in accordance with human musical convention (e.g. adherence to a human defined musical scale). Therefore, it produces a non-abstract sound signal representation of realistic protein interaction dynamics in which potential features of musicality can be investigated. Additionally, we develop a random forest model that uses 11 different features of the correlative structure of sound signals to accurately classify music, human speech, various animal vocalizations, and various nonbiological natural sounds. We then further examine feature importance with relation to these classifications and define feature shifts relative to music via a relative entropy-based divergence metric. This allows us to measure distances related to changes in a sound signals degree of musicality. In this study, we quantify the musicality of the AAV derived sounds of B-amyloid and various other protein interactions and compare them within the context of music and other sounds from the natural world.

**Figure 2A.**
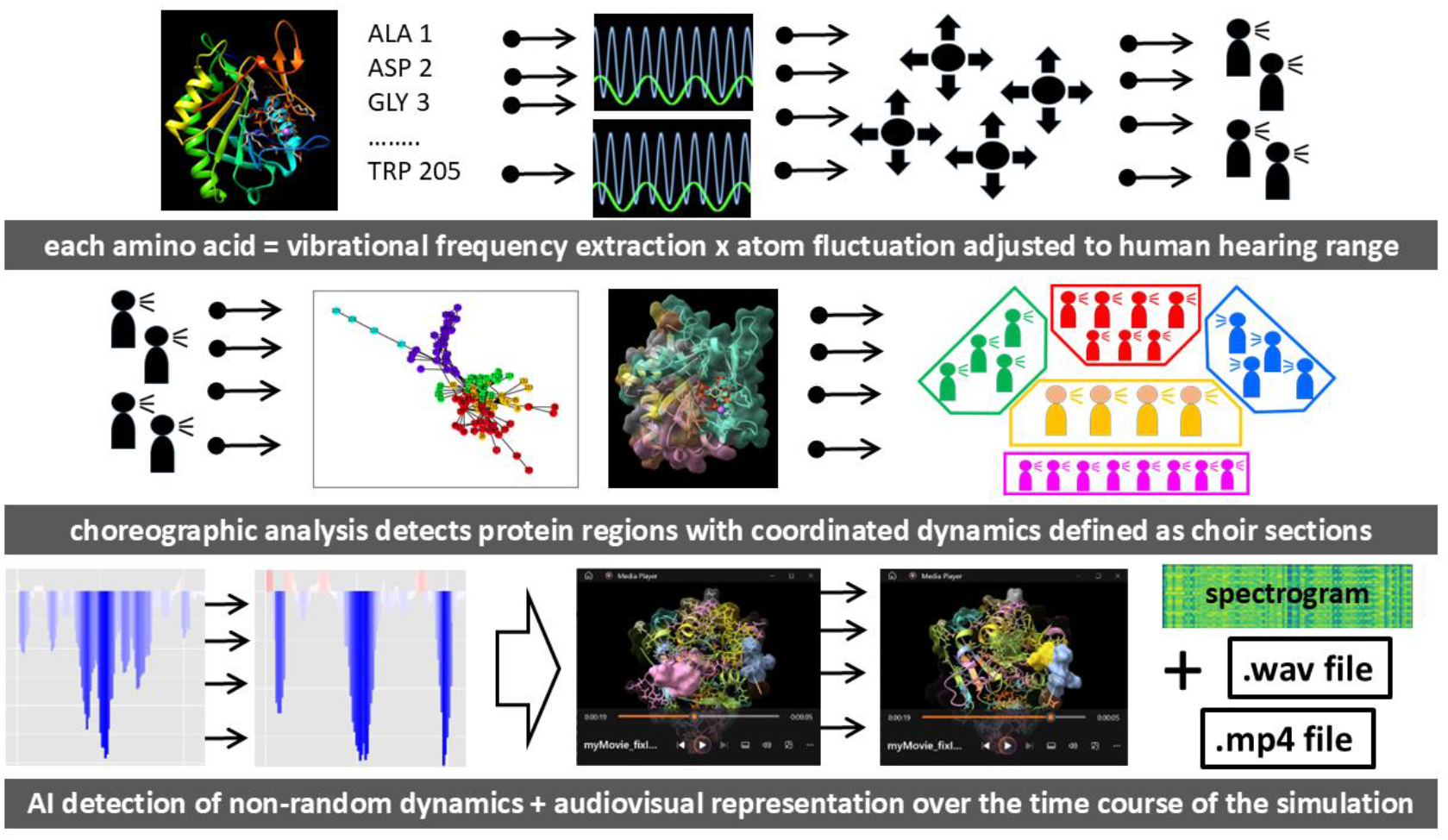
Graphical overview of the data sonification of protein interaction dynamics. Initially, vibrational frequencies are extracted per each amino acid and adjusted in accordance to average atom fluctuation and the pitch range of human hearing. Using choreographic analysis in ATOMDANCE, each amino acid frequency (i.e. individual voice) is layered in accordance to a community with which it shares coordinated dynamics (i.e. choir section). Using maxDemon denoising in ATOMDANCE, each frequency volume is adjusted according to its level of non-random shift in motion (i.e. functional binding) during protein interaction. Sound files and movie files are created by running this overall analysis pipeline over sub-segments of the molecular dynamic trajectory, subsequently creating each individual movie frame. See Supplemental File (AAV_examples.ppt) to see and hear examples of the this.

**Figure 2B.**
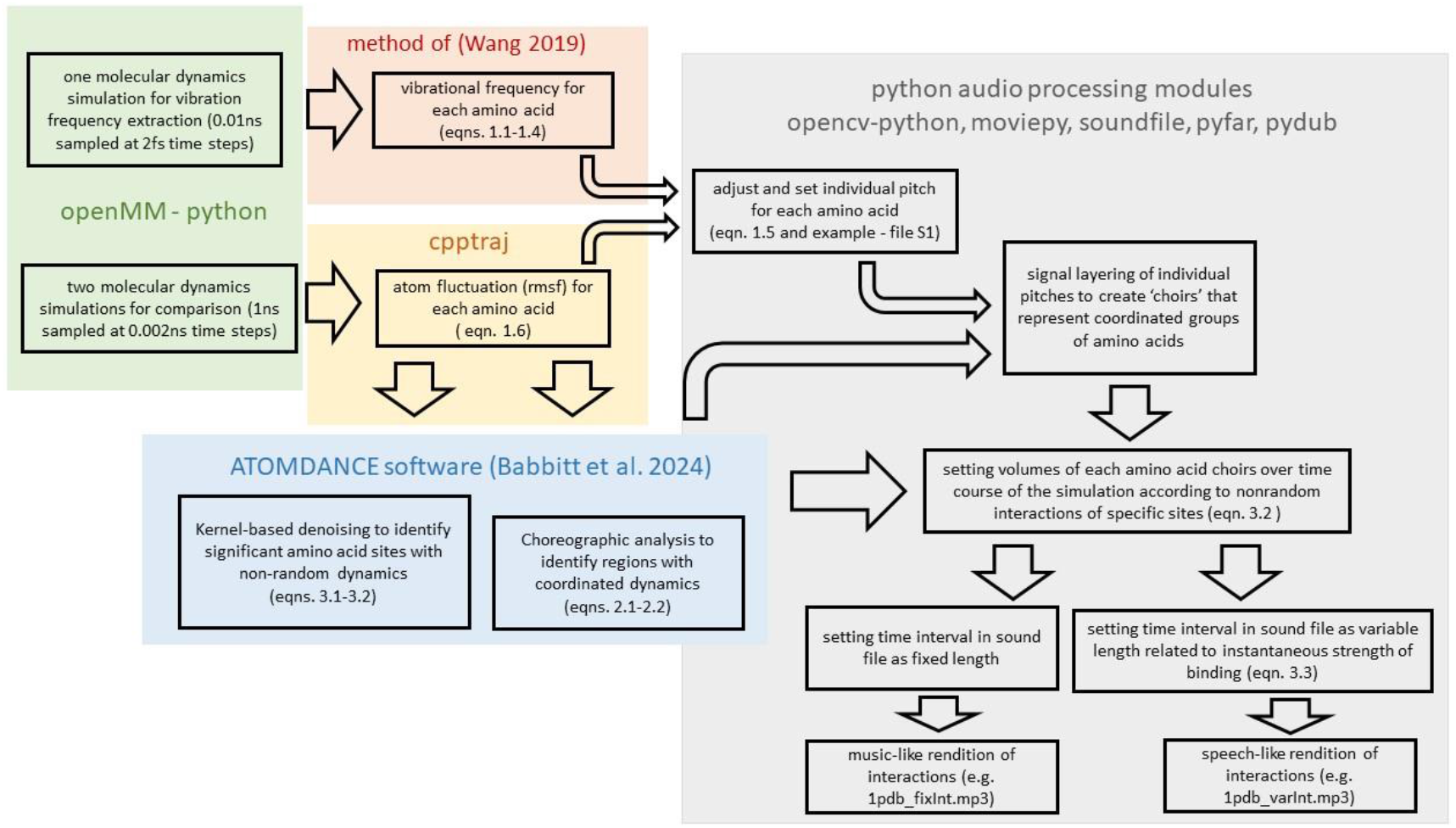
Schematic overview of the data sonification of protein interaction dynamics. All the methods and equations are fully defined in the Supplemental Methods section.

**Figure 2C.**
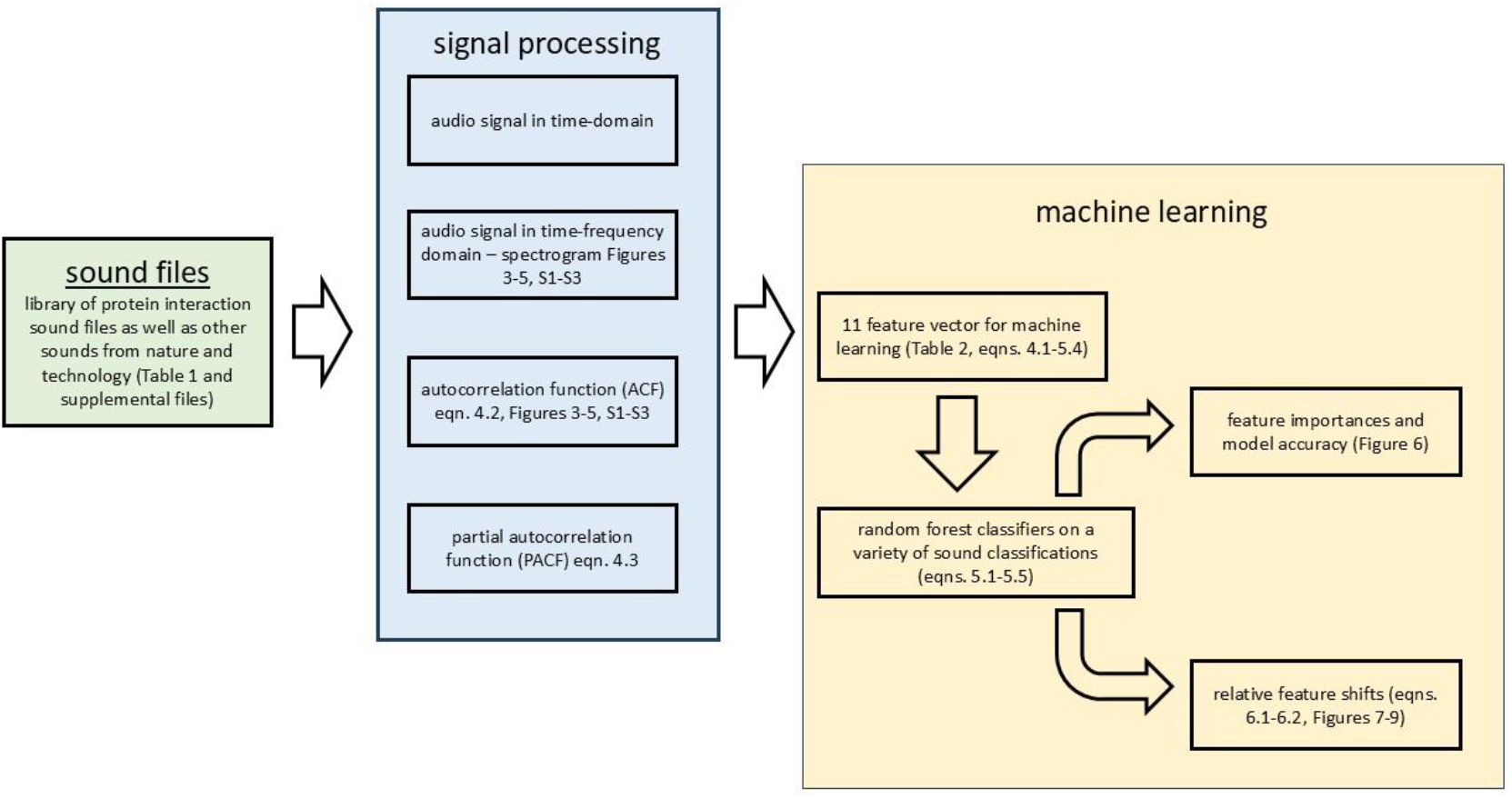
Schematic overview of the machine learning analysis of the sounds of protein interaction dynamics in the context of natural sounds. All the methods and equations are fully defined in the Supplemental Methods section.

## Results

### Spectral cross-correlation and time-domain autocorrelation

We find that AAV generated sounds of protein files exhibit sounds similar to that of a computer modem, but with more variability and complexity. AAV generated sounds with a fixed time interval are more musical sounding than those with a variable time interval adjusted to overall binding strength over time. However, their spectral and autocorrelative features are very similar, with fixed interval renditions appearing slightly less stationary (i.e. more triangular and less T shaped) in the autocorrelation function (ACF) plot (Figures 3, S1). Also, B-amyloid fibril interactions tend to have more complex ACF functions than do most other protein interactions (Figure 3). By comparison, spectrograms and ACF plots for a variety of technological and natural sounds (Figure 4), as well as different music (Figure 5) are also shown. Plots for computer modem, Morse code, and human crowd cheering are shown in 3A-F. As the signal becomes more layered (e.g. many voices in a large crowd), the ACF plot loses stationarity and becomes highly triangular. Plots for white, pink, and brown noise are also given in Figure 4D-F. Memory-less white noise (Figure 4D) has no significant peaks in the ACF as expected, and the number of distinct peaks in the ACF increases as the noise gains short-term memory (i.e. approaches brown noise) (Figure 4E-F). White noise also has extremely high note variability index (NVI) in addition to zero autocorrelation, whereas brown noise has extremely high first-order autocorrelation and lower NVI. The ACF plots for music exhibit a wide variety of ACF behavior, ranging from more stationary signals of classical music (Figure 5A-B) to more layered stationary signals of modern minimalist electronic music (Figure 5C-E) to far more layered nonstationary signals in modern electronic music (Figure 4F). Spectrograms and ACF plots for human speech and a variety of animal vocalizations (Figure S2), and non-biological natural sounds (Figure S3) are also shown. Human speech is characterized by high stationarity and little signal layering in the ACF when compared to most other animal vocalizations (Figure S2A).

**Figure 3.**
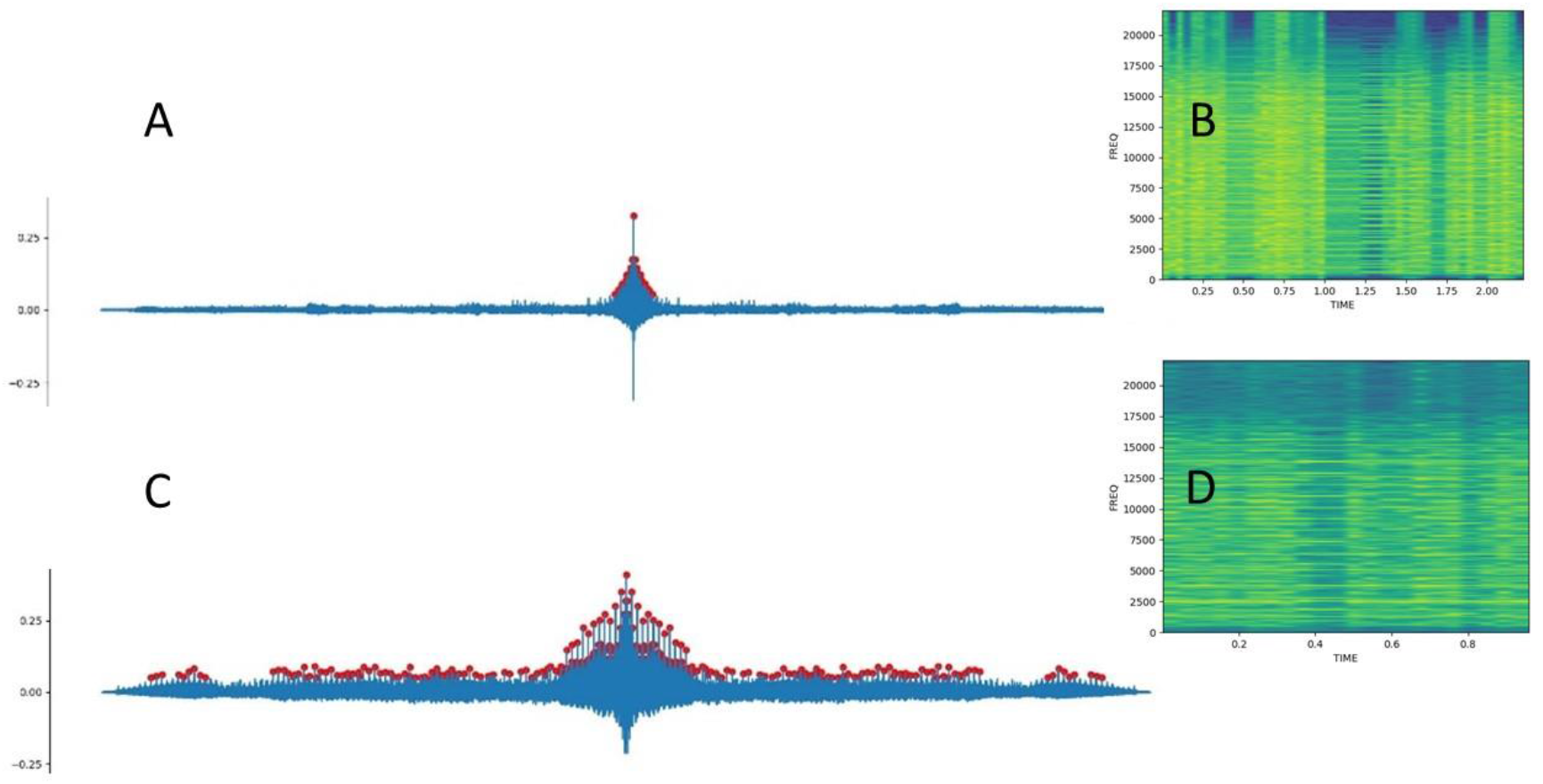
The autocorrelation plots (A,C) and time-frequency domain spectrograms (B,D) for the interaction of TATA binding protein with DNA (A,B) and amyloid fibril interactions (C, D) using variable time intervals during the analysis where the length of interval represents the overall strength of binding during each time frame of the analysis. Here a shorter time interval represents stronger binding at that given point in time. Distinct peaks in the autocorrelation function are shown in red.

**Figure 4.**
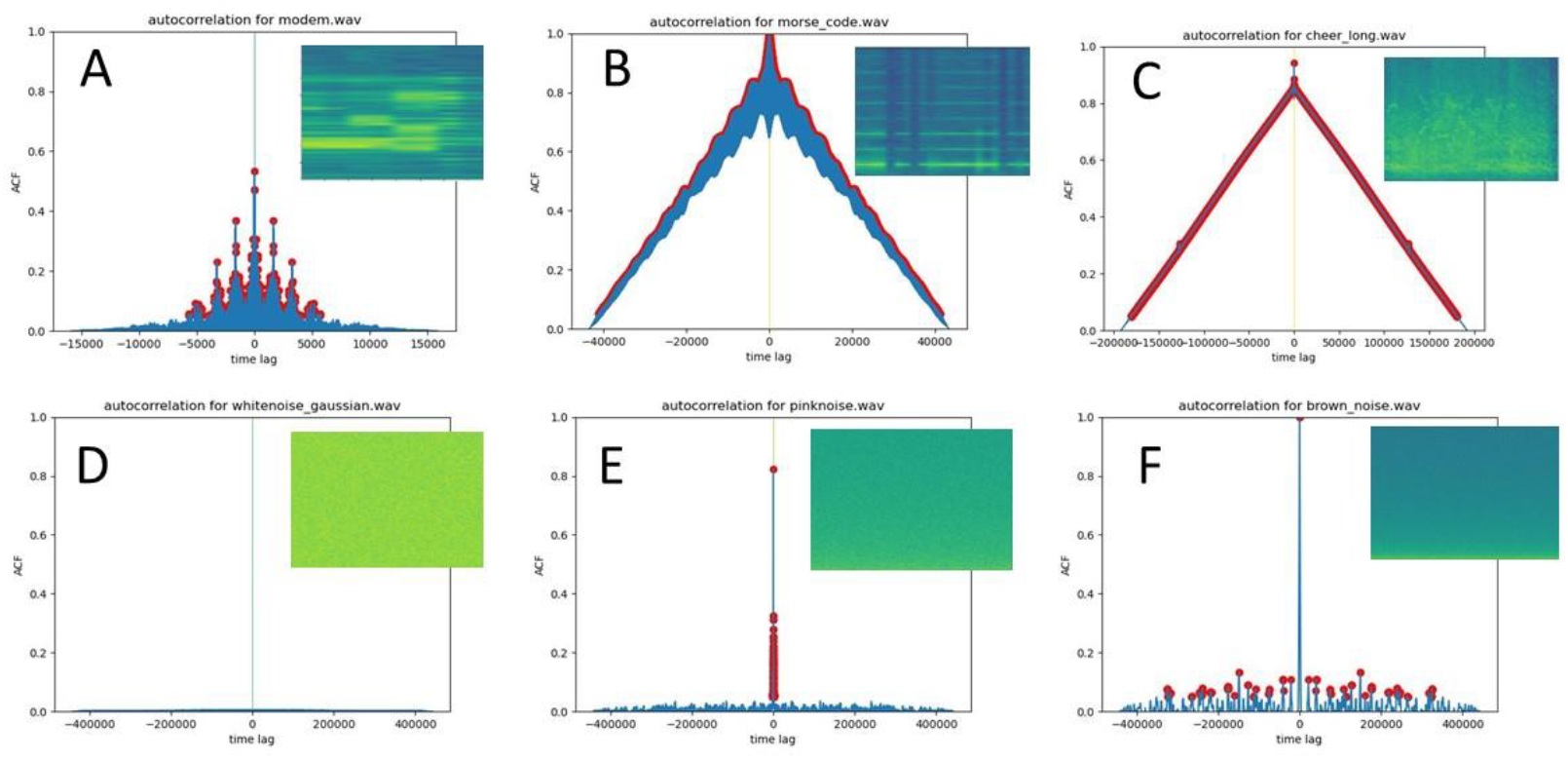
Autocorrelation plots and spectrograms (inset) for technological and natural sounds of (A) computer modem, (B) morse code, (C) a large crowd cheering at a game, (D) memory-less Gaussian white noise, (E) power law distributed pink noise and (F) Brownian noise (i.e. random walk with memory). Distinct peaks in the autocorrelation function are shown in red.

**Figure 5.**
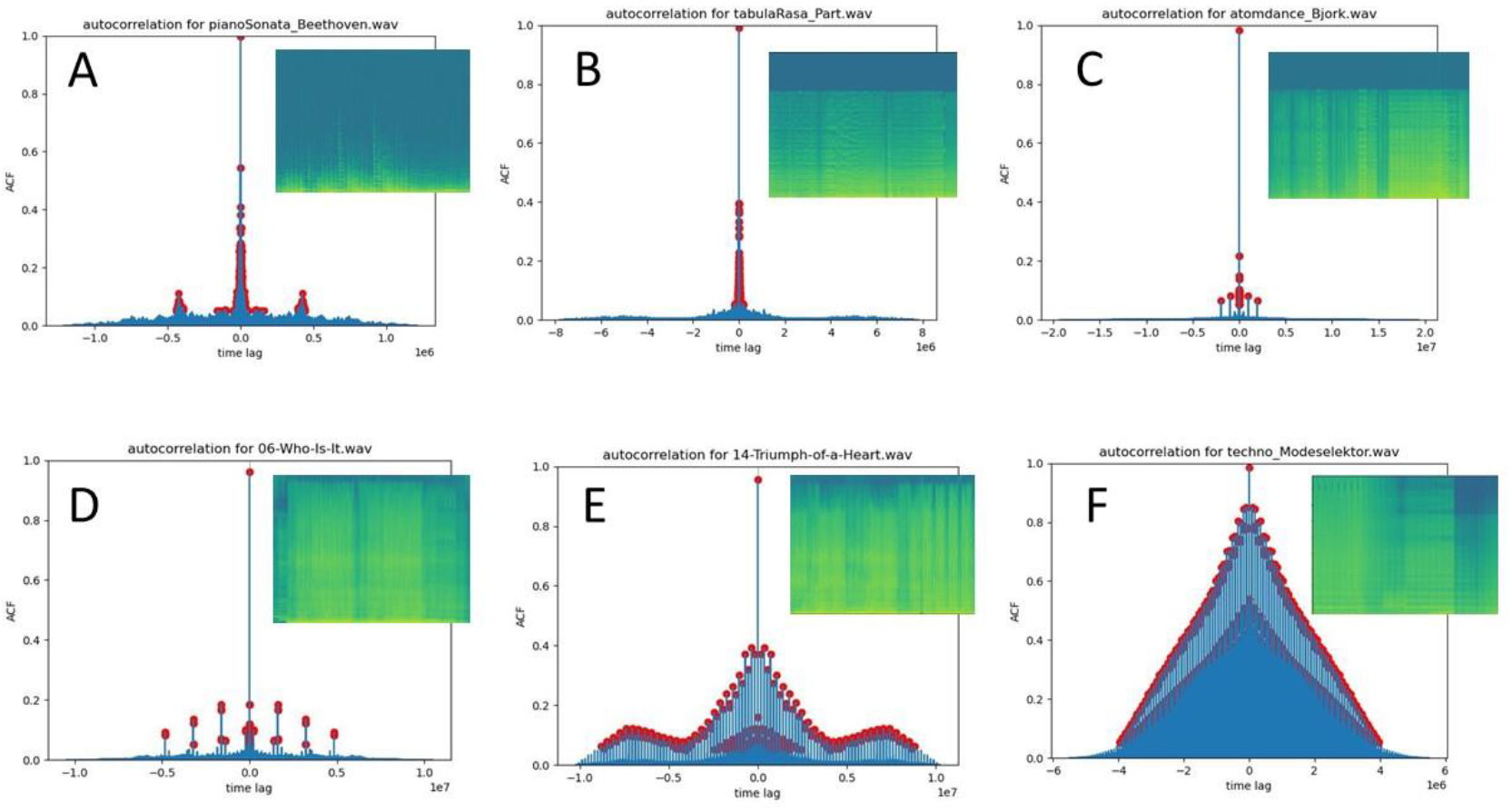
Autocorrelation plots and spectrograms (inset) for examples of human music including a (A) piano sonata by Ludwig Van Beethoven, (B) Tabula Rasa by the classical minimalist Arvo Part, (C) Atom Dance, a minimalist piece by Björk (D) Who Is It, a vocals-only pop song by Björk, (E) Triumph of a Heart, a vocals-only pop song by Björk and (F) Ohm, a hardcore techno piece by Modeselektor. Distinct peaks in the autocorrelation function are shown in red.

All .wav and .mp3 sound files, and plots for time-domain, ACF, PACF, and spectrogram for all sounds analyzed in this study are given in Supplementary Data (DOI: 10.5281/zenodo.15013666). Some illustrative examples are shown as a slide show in the supplemental file AAV_examples.ppt equipped with embedded mp3 and mp4 files.

### Random forest classification of music, vocalization, and non-biological sound

The various feature importances for binary classifications comparing different kinds of sounds is shown (Figure 6A). In summary, first order AC is most important in distinguishing protein interactions for other sounds, the number of ACF peaks is most important for distinguishing music from human speech, and NVI is most important for distinguishing nonbiological sounds from biological sounds including music. Almost all binary comparisons are highly accurate (i.e. > 95%) for almost all comparisons involving music, human and animal vocalizations, and other natural sounds (Figure 4B). Several exceptions are an inability to discriminate ocean waves from flowing water, and reduced accuracy in discriminating between animal vocalizations of major taxonomic groups and between several music genres. However, the main results of this study did not depend upon these weaker classifications. Bar plots for feature importances and boxplots for comparative feature values for all comparisons in Figure 4 are given in Supplementary Data (DOI: 10.5281/zenodo.15013666).

**Figure 6.**
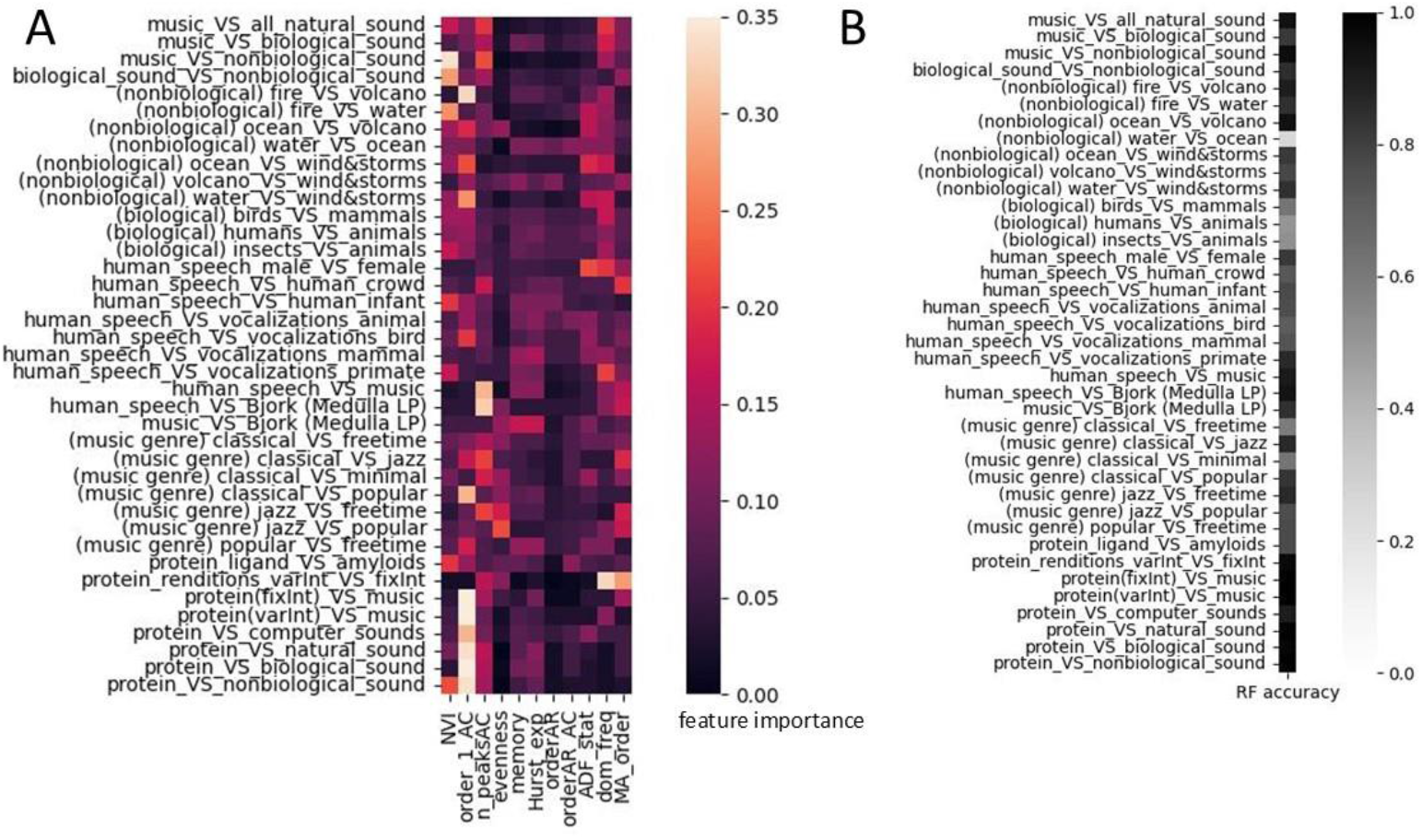
A random forest classifier trained on the 11 acoustic features for discerning musical from nonmusical sounds. Heatmaps showing (A) feature importance, and (B) overall accuracy are shown for each binary classification (i.e. vertical axes). The accuracy for most classifications discerning music and speech from other biological and non-biological sounds are very high (> 90%).

### Acoustic feature shifts in music and human vocalization

Acoustic feature shifts (AFS) determined via differences in statistical divergence (i.e. JS distances) relative to music are very interesting. We report that the AAV generated sounds of highly structurally conforming B-amyloid fibril interactions are indeed more musical (i.e. negative shifted) and more speech-like than other non-conforming protein interactions (Figure 7). This can also be readily heard in the several sound file example also provided (Supplemental File S1). While the overall shift is relatively small (−5.5%) and only slightly discernable to the ear, in both cases it is driven by a significant 24% shift in first order autocorrelation and a less significant 10-12% shift in signal persistence or memory level. We also demonstrate the female speech is also more musical than male speech, exhibiting positive shifts in note variability (i.e. complexity) and negative shifts in dominant frequency, order of the moving average and the r value of the correlation observed at the order of the autoregression function (Figure S5). When we compared human speech to music in reference to other animal vocalizations, we find that human speech is shifted farther away from other animal vocalizations than is human music, due largely to the r value at the order of the autoregression Figure 8. As music is quite different from human speech in the way it is generated, often involving various forms of instrumentation, we also did this analysis using 14 tracks from the Icelandic artist Bjork’s 5^th^ album as a control (8A-B). In this work, the artist strived to replicate her own modern musical style without using any instrumentation but instead using purely vocal techniques of the beatboxer Razhel, Inuit throat singer Tanya Tagaq and a small Greenlandic choir to replicate missing instrumentation. We find that this control (8A-B) produced nearly identical results regarding acoustic feature shift when compared to the same analysis using all the music files in our database (Figure 8C-D). This indicates that human speech is more acoustically derived from animal vocalization than is music, regardless of the effect of instrumentation, suggesting that it is likely more evolutionarily derived as well (i.e. musicality in human voice predated the evolution of speech). Lastly, we compared male and female speech in reference to both the vocalizations of animals (Figure 9A-B) and human infants (Figure 9C-D). Here we find that male speech is more acoustically derived than female speech, perhaps suggestive of greater natural selection on the acoustic features of the male voice.

**Figure 7.**
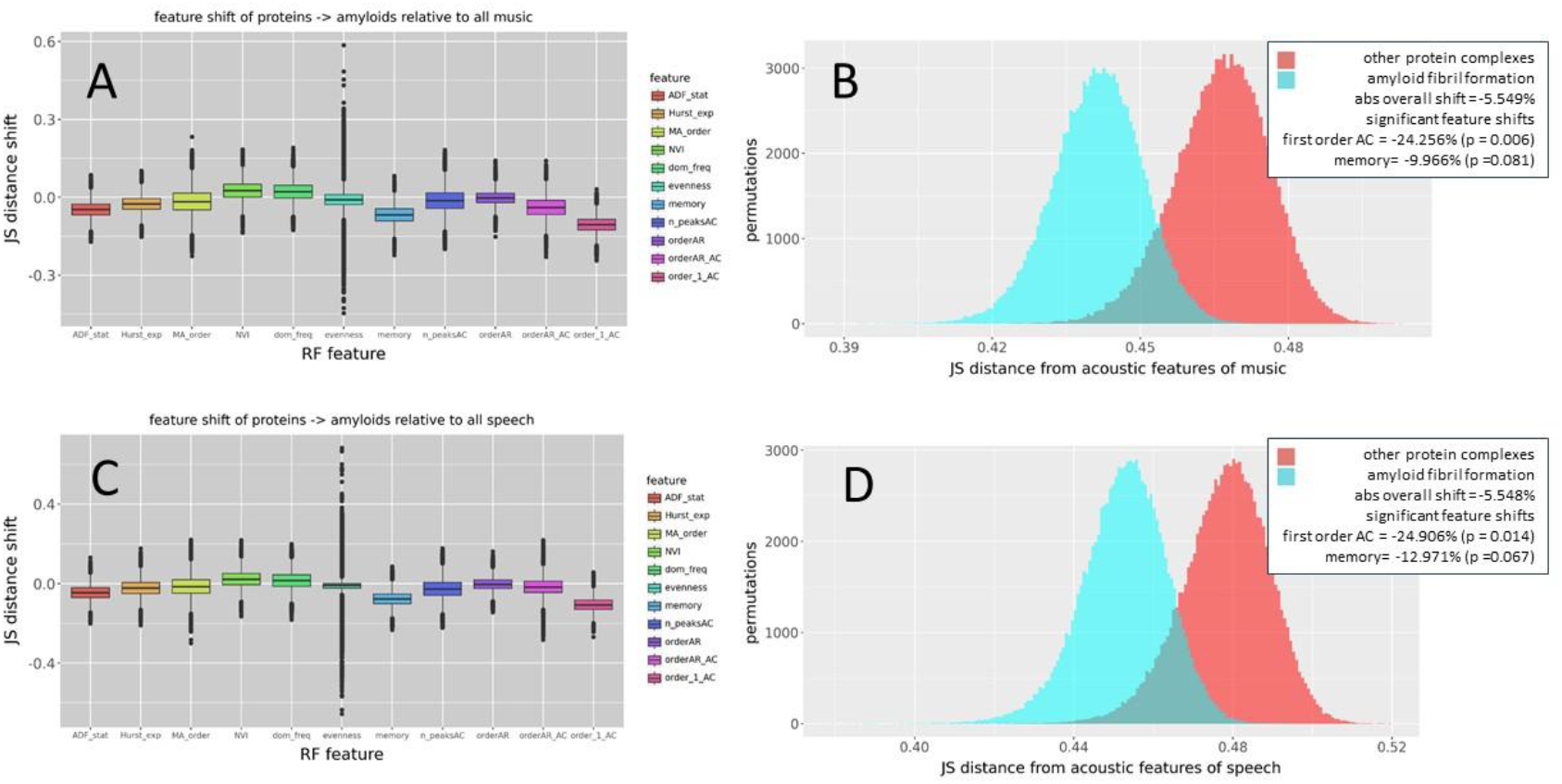
Acoustic feature shifts of data sonification of interaction dynamics of amyloid fibrils compared to other proteins and other proteins in reference to (A-B) music and (C-D) human speech. The distribution shifts are calculated as differences in Jensen-Shannon distance between bootstrapped sample distributions. Note: positive shift values in A and C indicate movement away from the reference distribution whereas negative shifts indicate movement towards it. Empirical p-values for each feature are determined by permutations tests (n = 100000).

**Figure 8.**
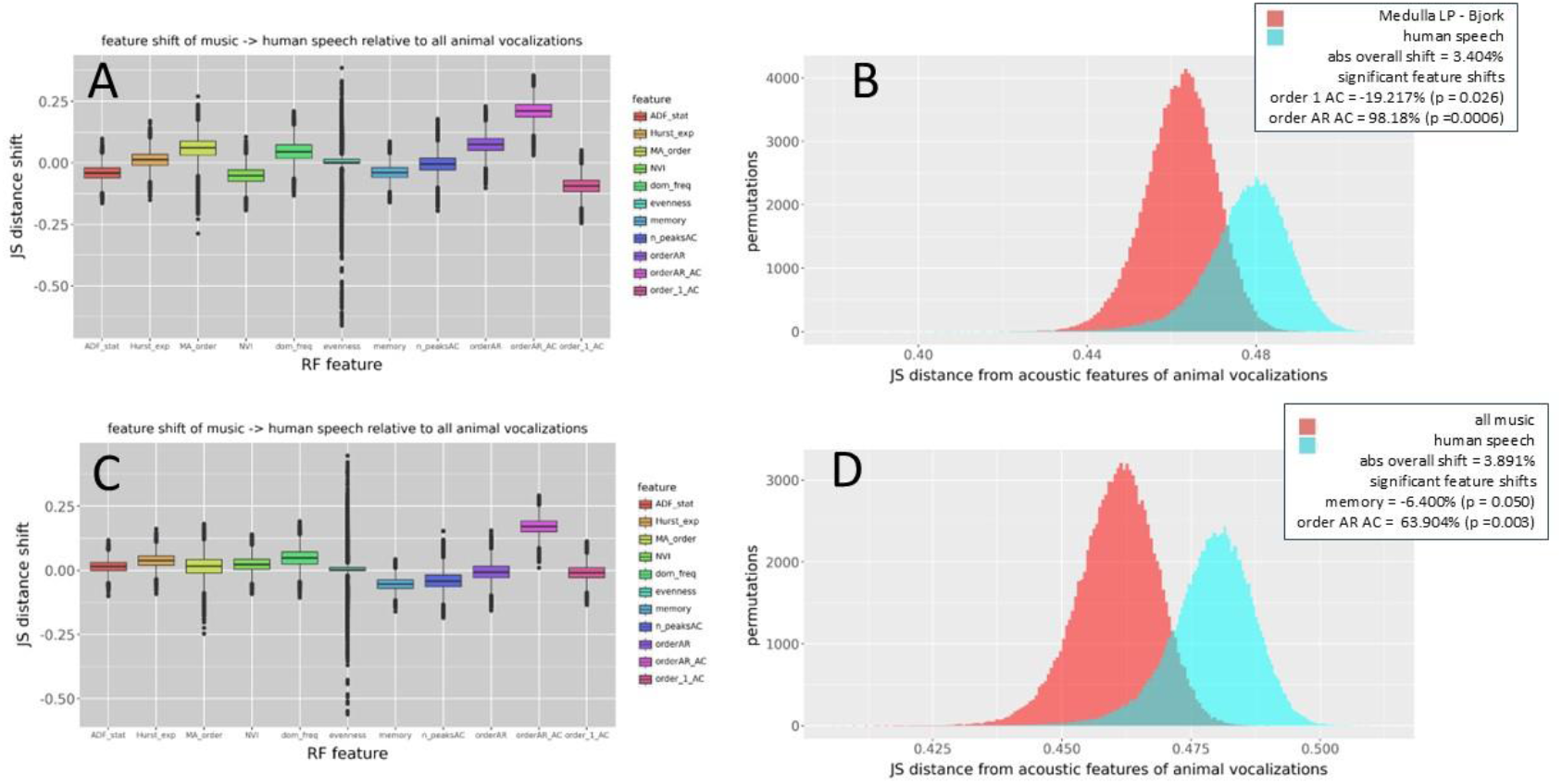
Acoustic feature shifts for human speech compared to music in reference to animal vocalization. For the representations of music we use both (A-B) Björk’s 4^th^ solo album Medulla where instrumentation was replaced nearly entirely by human voice generated sounds and (C-D) a much larger catalog of music with various forms of instrumentation. The distribution shifts are calculated as differences in Jensen-Shannon distance between bootstrapped sample distributions. Note: positive shift values in A and C indicate movement away from the reference distribution whereas negative shifts indicate movement towards it. Empirical p-values for each feature are determined by permutations tests (n = 100000).

**Figure 9.**
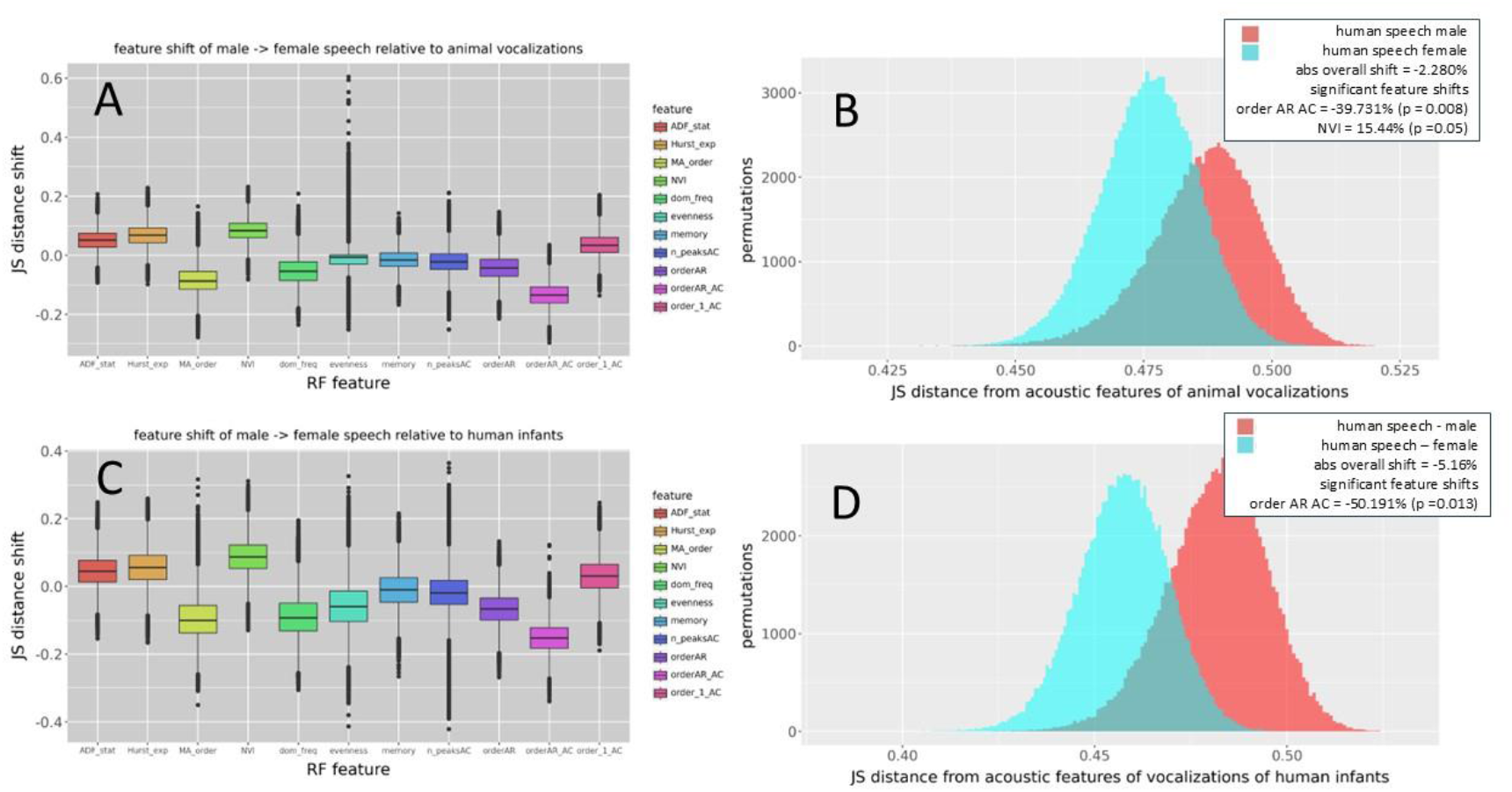
Acoustic feature shifts for human speech (female) compared to human speech (male) in relation to animal vocalizations (A-B) and the sounds of human infants (C-D). The distribution shifts are calculated as differences in Jensen-Shannon distance between bootstrapped sample distributions. Note: positive shift values in A and C indicate movement away from the reference distribution whereas negative shifts indicate movement towards it. Empirical p-values for each feature are determined by permutations tests (n = 100000).

## Discussion

In this study, we have developed a non-abstract method of data sonification applied to physics-based computer simulations of dynamic protein interactions that we can employ to analyze resulting sounds for features of musicality. As musical sound is highly autocorrelated when compared to other sounds, we developed a set of 11 correlative mathematical features (i.e. feature vector) that allows for highly accurate machine learning discrimination of human speech and music from a variety of other biological and non-biological natural sounds. This random forest model embeds feature importance as decision tree depth, allowing us to observe what individual features are important in discriminating various natural sounds including those created by our AAV pipeline. Through the analysis of comparative shifts in acoustic features relative to features defining music, we demonstrate that protein complexes comprised of components that are structurally conforming (i.e. amyloids) are more speech-like and musical in their resulting sounds. We hypothesize that musicality may play a general role in social behavioral conformity at multiple system scales, including human prosocial evolution, for which there is previous evidence from a variety of other fields (neurophysiology, biomusicology, and acoustic archeology). Analysis of feature shifts in our data with respect to animal vocalization reveals that music is less acoustically derived from animal vocalization than human speech. This holds even when controlling for the use of instrumentation in music. As animal vocalizations are an evolutionary ground point for human music and language, the finding that speech is more derived than music suggests that the evolution of musicality in the human voice may have predated its use in language. We also find that sex differences in feature shifts of human vocalizations relative to other animals and human infants suggests that the acoustic characteristic of the male voice is more derived than that of females, perhaps also indicating differences in voice arising through a history of stronger sexual selection acting on the male voice.

Our work has several important strengths and weaknesses. While we find that overall feature shifts are small (5%), we find that within specific features they can be quite large (>20%). When analyzing amyloid interaction, we found the shift in first order autocorrelation (i.e. ambience or dimension) and signal memory (i.e. persistence) towards musicality are both particularly strong. We think this result speaks to the importance of molecular resonance in the characteristic solenoid structures of amyloids in replicating the resonance of musical sounds in particular instruments and ambient spaces in which we tend to enjoy producing music (i.e. concert halls, recording studios, bathrooms, subway tubes, caves, and canyon walls). Another major strength of our work is the objectivity provided using machine learning in place of human discrimination as well as the use of a non-abstract means of data sonification rather than human musical conventions when producing protein sounds to analyze and compare. However, our work is potentially limited by the relatively short simulation times needed to compensate for the computational cost of the whole study. Short simulations may have prevented the manifestation of some of the long-term correlative aspects in sound signals, if they were present.

In recent years, there has been a lot of interest in defining the evolutionary origin and purpose of music. It is relatively clear that the role of music in modern society shares much of the same primordial goals of other vocalizations and displays in the animal world. The often-sexualized content of modern popular music may reflect a common origin to animal vocalization in its function to sexually attract potential mates, display individual fitness via acts of physical performance, and to organize physical and emotional states of larger groups. When this latter aspect of music is related to our tendency towards prosocial behavior, it can perhaps also explain why music gives us feelings of transcendence as well. In transcendent states, we find release from our focus as individuals, becoming a smaller part of a larger and more organized group existence. While much about music is subjective and cultural, we conclude that the correlative or co-relational acoustic features of our natural soundscapes are likely key common ancestral neurophysiological signals in defining the boundaries of music, language, and non-human animal vocalizations relative to other sounds in nature. While music itself is typically defined as a product of human culture, our results also strongly suggest that certain aspects of musical sounds (i.e. musicality) can be used to quantitatively define acoustic metrics that may defy the boundary of human uniqueness. Along with recent discovery of elements of musicality in other animals, this is highly suggestive of broader evolutionary function for a general type of signal we describe as ‘musical’. As music is highly autocorrelated, we conjecture that a proximate evolutionary function of musicality is to help predict and instruct group behavior in social settings, allowing groups to achieve common goals (i.e. achieve prosocial behavior). Because prosocial behavior can be extended to certain highly conforming protein complexes such as amyloids, we suggest that musicality may also be a physical feature of the natural world that extends across many scales in biology. This might include the domains of the molecular, cellular, organismal, and perhaps even the ecological. The idea that music may have a physical role in nature apart from humanity is not new and it is sometimes even a creative point of view of music itself (e.g. Pythagoras and Kepler’s concept of harmony/music of the spheres, Björk’s biophilia project). In the future, it should be interesting to apply more advanced machine learning methods to better determine what elements of musicality fall within versus beyond human experience.

## Supporting information

examples in .ppt

## Acknowledgements

We’d like to acknowledge the many conversations we have had with faculty colleagues, fellow musicians, and former and current students in many disciplines across the arts and sciences. We are particularly grateful for the comments of Erica Haskell and Joel Hunt (RIT School of Performing Arts) and Joaquin Carbonara (Computer Science at Buffalo State University) during the early stages of this research. We thank Kiersten Winter for her work on the early stages of the code. We are grateful to the open-source software community for producing many of the software tools utilized in this research. We are thankful for the many fascinating questions and points of view put forth in the 2011 short film “When Björk met Attenborough” including those of luminaries Sir David Attenborough, Björk Gudmundsdottir, Oliver Sacks, and their many collaborators. This research was not supported by Federal grants.

---

**Figure S1.**
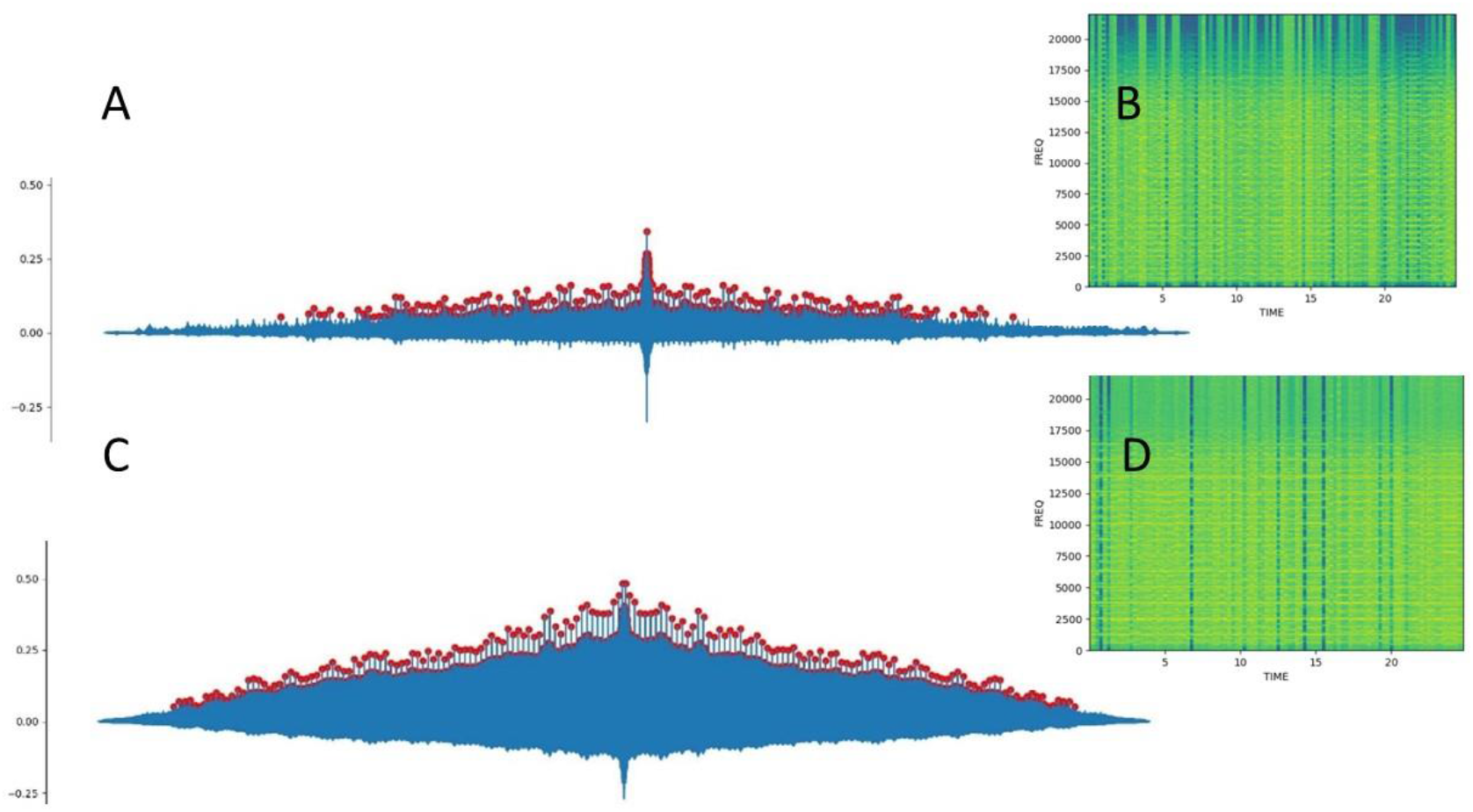
The autocorrelation plots (A,C) and time-frequency domain spectrograms (B,D) for the interaction of TATA binding protein with DNA (A,B) and amyloid fibril interactions (C, D) using fixed time intervals during the analysis. Distinct peaks in the autocorrelation function are shown in red. NOTE: The sonic rendition of protein interactions with fixed intervals of time yields a more musical sound with a constant cadence than renditions with variable length intervals (i.e. compare to Figure 3).

**Figure S2.**
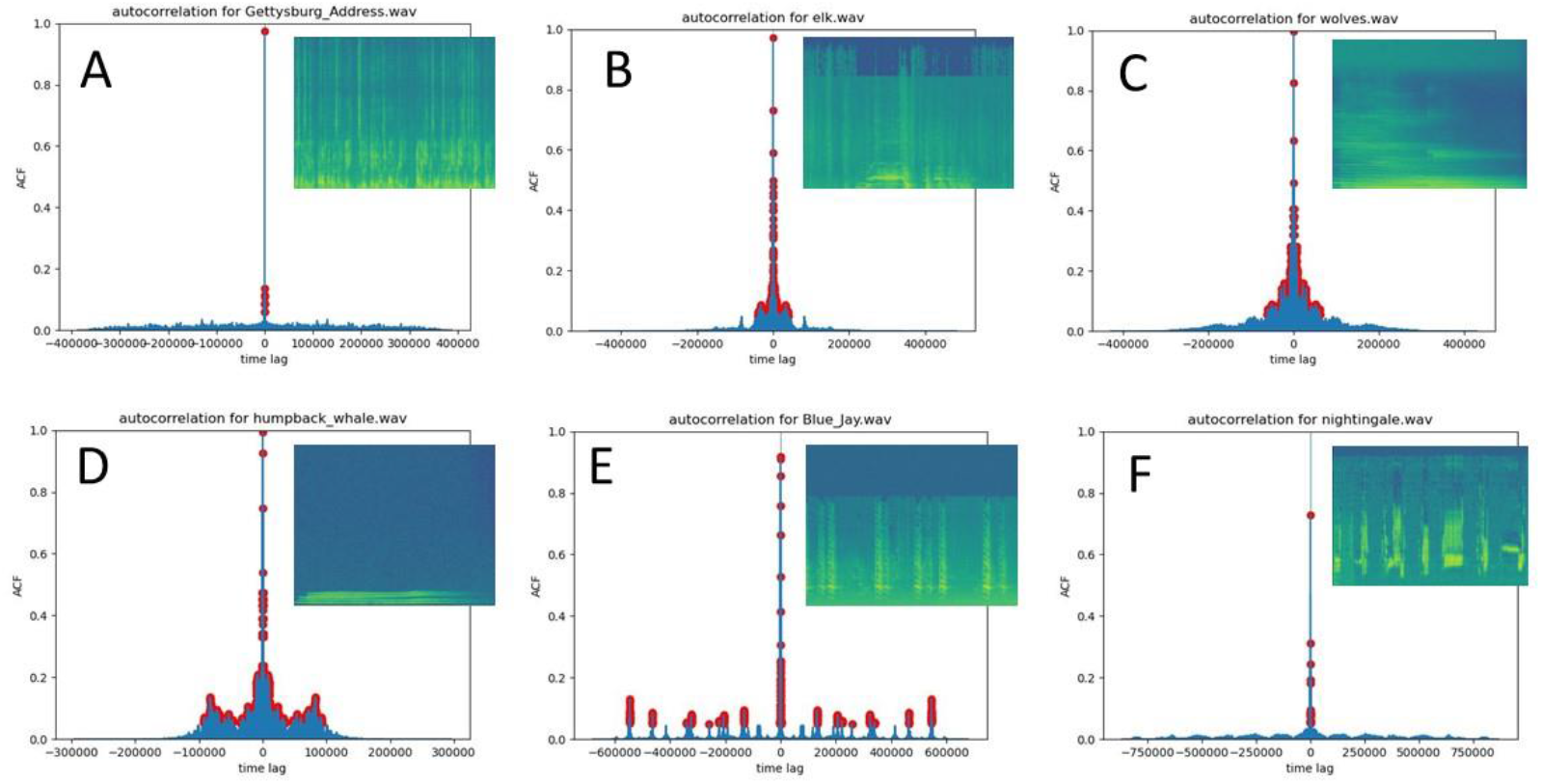
Autocorrelation plots and spectrograms (inset) for human and animal vocalizations of a (A) single human speaking, (B) a single elk bugling, (C) wolves howling (D) humpback whales singing, (E) a Blue Jay calling and (F) a Nightingale singing. Distinct peaks in the autocorrelation function are shown in red.

**Figure S3.**
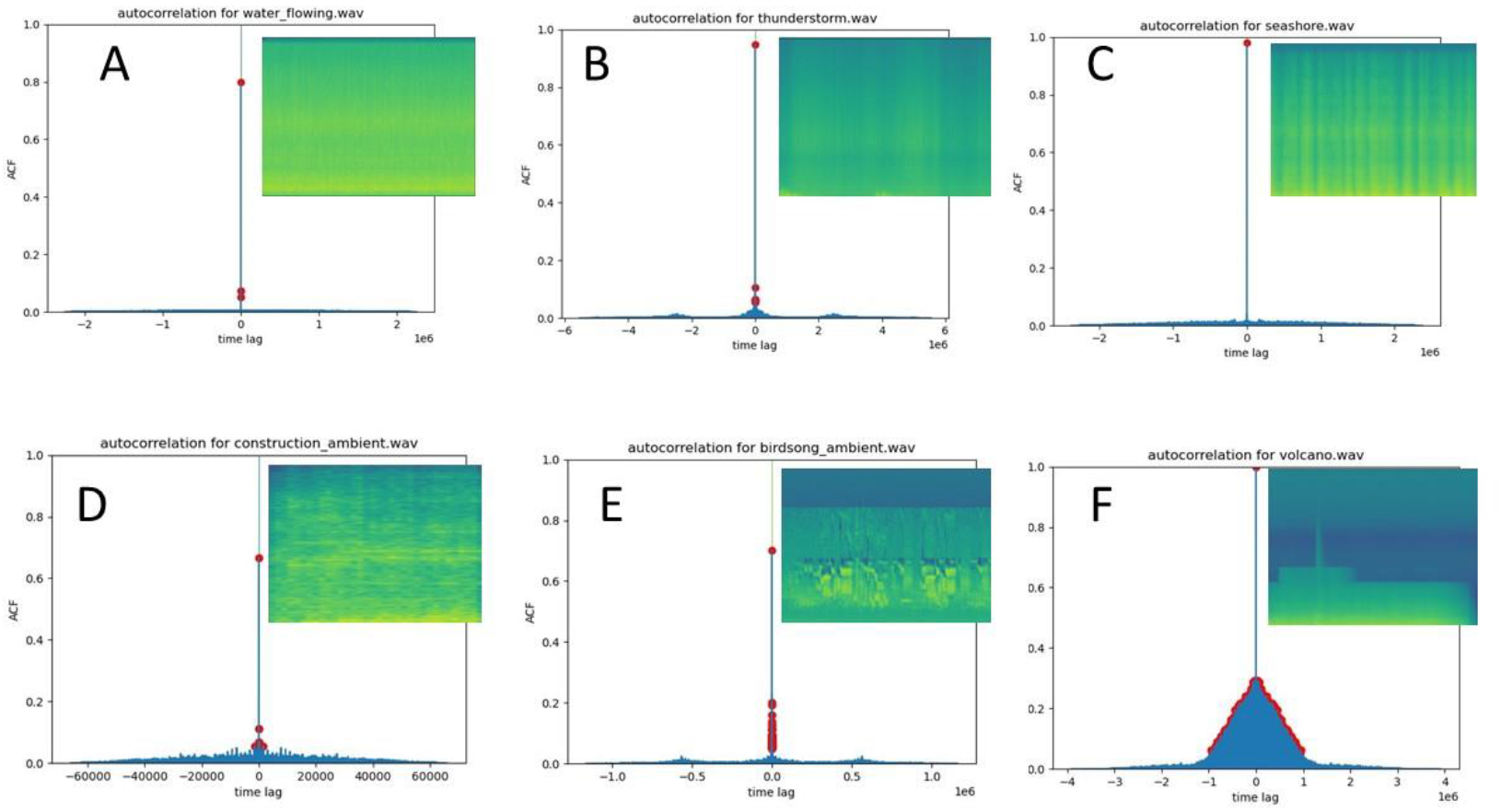
Autocorrelation plots and spectrograms (inset) for natural sounds of (A) water flowing in a stream, (B) a thunderstorm, (C) an ocean seashore (D) a construction site, (E) a forest with bird song, and (F) a volcano erupting. Distinct peaks in the autocorrelation function are shown in red.

**Figure S4.**
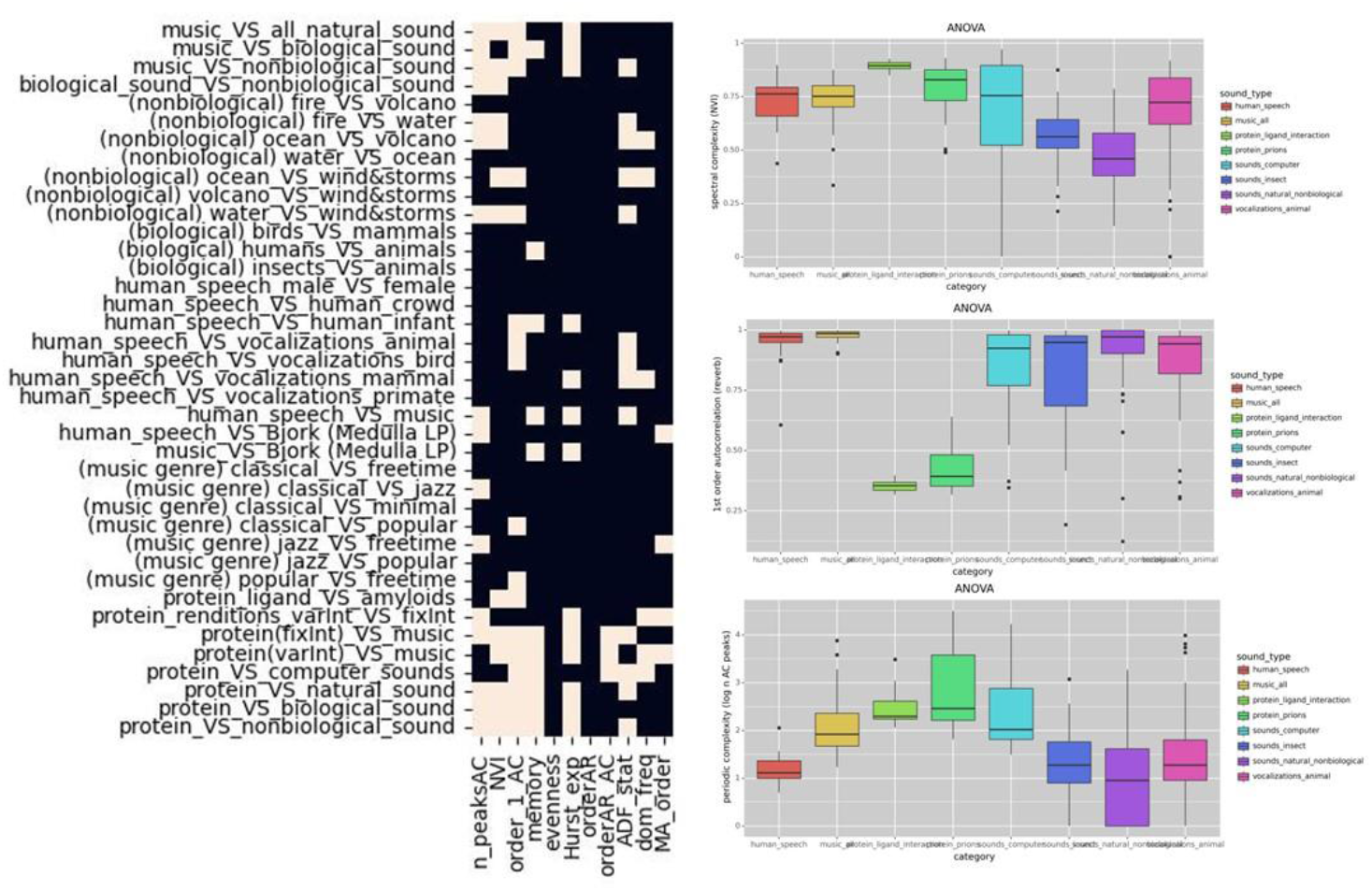
Heatmaps for significance of t-test on each acoustic feature (left) and some examples of boxplots comparing major sound categories on the three most important acoustic features in the random forest model (right).

**Figure S5.**
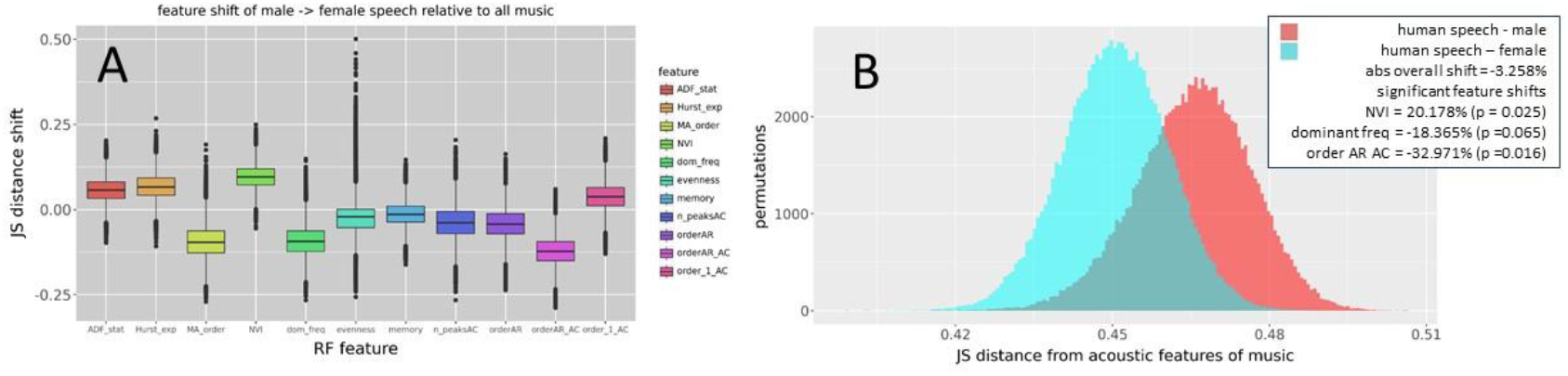
Acoustic feature shifts of male and female speech in reference to music. The distribution shifts are calculated as differences in Jensen-Shannon distance between bootstrapped sample distributions. Note: positive shift values in A indicate movement away from the reference distribution whereas negative shifts indicate movement towards it. Empirical p-values for each feature are determined by permutations tests (n = 100000).

**Figure S6.**
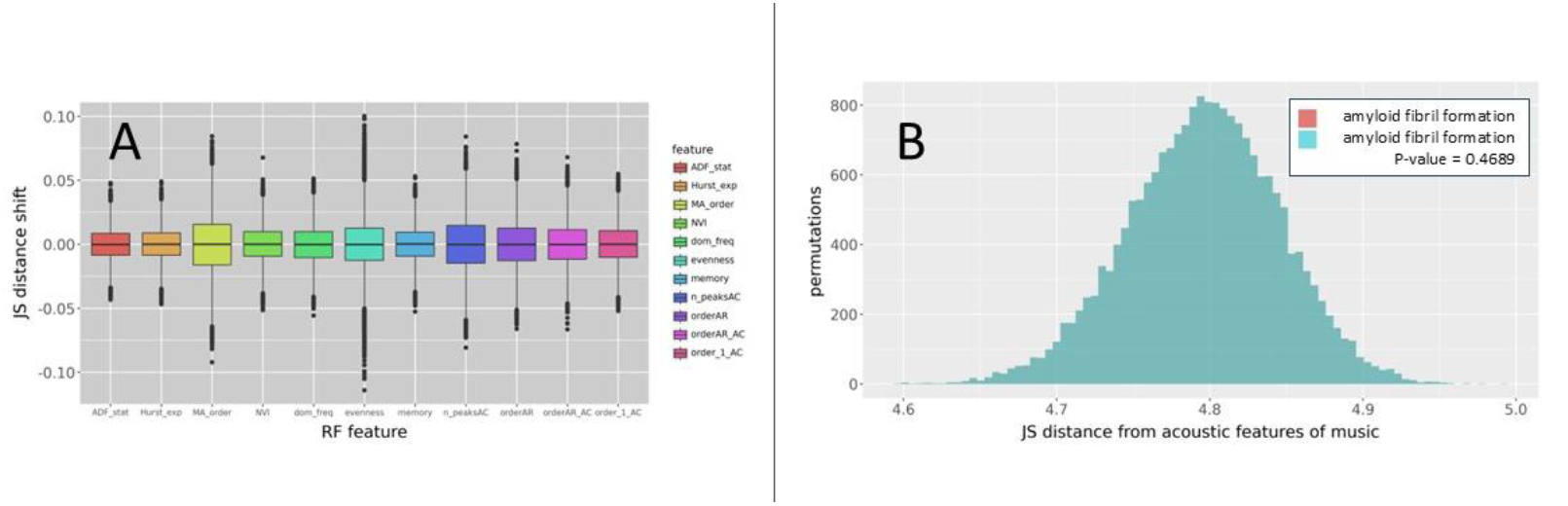
Null hypothesis generated test for our method of quantifying acoustic feature shift (Figures 7-9). Here we use the same data distribution for P1, P2 and Q in the Jensen-Shannon equations.

## References

Babbitt, G.A., Rajendran, M., Lynch, M.L., Asare-Bediako, R., Mouli, L.T., Ryan, C.J., Srivastava, H., Rynkiewicz, P., Phadke, K., Reed, M.L., Moore, N., Ferran, M.C., Fokoue, E.P., 2024. ATOMDANCE: Kernel-based denoising and choreographic analysis for protein dynamic comparison. Biophys. J. 123, 2705–2715. 10.1016/j.bpj.2024.03.024

Baptista, L.F., Keister, R.A., 2005. Why birdsong is sometimes like music. Perspect. Biol. Med. 48, 426–443. 10.1353/pbm.2005.0066

Bedoya, D., Arias, P., Rachman, L., Liuni, M., Canonne, C., Goupil, L., Aucouturier, J.-J., 2021. Even violins can cry: specifically vocal emotional behaviours also drive the perception of emotions in non-vocal music. Philos. Trans. R. Soc. B Biol. Sci. 376, 20200396. 10.1098/rstb.2020.0396

Bilger, H.T., Vertosick, E., Vickers, A., Kaczmarek, K., Prum, R.O., 2021. Higher-Order Musical Temporal Structure in Bird Song. Front. Psychol. 12. 10.3389/fpsyg.2021.629456

Brown, S., Savage, P.E., Ko, A.M.-S., Stoneking, M., Ko, Y.-C., Loo, J.-H., Trejaut, J.A., 2014. Correlations in the population structure of music, genes and language. Proc. R. Soc. B Biol. Sci. 281, 20132072. 10.1098/rspb.2013.2072

Chang, A., Teng, X., Assaneo, M.F., Poeppel, D., 2024. The human auditory system uses amplitude modulation to distinguish music from speech. PLOS Biol. 22, e3002631. 10.1371/journal.pbio.3002631

Chatani, E., Yuzu, K., Ohhashi, Y., Goto, Y., 2021. Current Understanding of the Structure, Stability and Dynamic Properties of Amyloid Fibrils. Int. J. Mol. Sci. 22, 4349. 10.3390/ijms22094349

Crespi, B.J., Yanega, D., 1995. The definition of eusociality. Behav. Ecol. 6, 109–115. 10.1093/beheco/6.1.109

De Gregorio, C., Maiolini, M., Raimondi, T., Carugati, F., Miaretsoa, L., Valente, D., Torti, V., Giacoma, C., Ravignani, A., Gamba, M., 2024. Isochrony as ancestral condition to call and song in a primate. Ann. N. Y. Acad. Sci. 1537, 41–50. 10.1111/nyas.15151

Fitch, W.T., 2015. Four principles of bio-musicology. Philos. Trans. R. Soc. Lond. B. Biol. Sci. 370, 20140091. 10.1098/rstb.2014.0091

Fitch, W.T., 2006. The biology and evolution of music: a comparative perspective. Cognition 100, 173–215. 10.1016/j.cognition.2005.11.009

Gallese, V., Gernsbacher, M.A., Heyes, C., Hickok, G., Iacoboni, M., 2011. Mirror Neuron Forum. Perspect. Psychol. Sci. J. Assoc. Psychol. Sci. 6, 369–407. 10.1177/1745691611413392

Galton, F., 1888. Co-Relations and Their Measurement, Chiefly from Anthropometric Data. Proc. R. Soc. Lond. 45, 135–145.

Gordon, C.L., Cobb, P.R., Balasubramaniam, R., 2018. Recruitment of the motor system during music listening: An ALE meta-analysis of fMRI data. PLoS ONE 13, e0207213. 10.1371/journal.pone.0207213

Honing, H., 2018. On the biological basis of musicality. Ann. N. Y. Acad. Sci. 10.1111/nyas.13638

Honing, H., ten Cate, C., Peretz, I., Trehub, S.E., 2015. Without it no music: cognition, biology and evolution of musicality. Philos. Trans. R. Soc. B Biol. Sci. 370, 20140088. 10.1098/rstb.2014.0088

Hooper, P.L., Gurven, M., Winking, J., Kaplan, H.S., 2015. Inclusive fitness and differential productivity across the life course determine intergenerational transfers in a small-scale human society. Proc. R. Soc. B Biol. Sci. 282, 20142808. 10.1098/rspb.2014.2808

Izbicki, P., Zaman, A., Stegemöller, E.L., 2020. Music Form but Not Music Experience Modulates Motor Cortical Activity in Response to Novel Music. Front. Hum. Neurosci. 14, 127. 10.3389/fnhum.2020.00127

Jensen, H.J., 1998. Self-Organized Criticality: Emergent Complex Behavior in Physical and Biological Systems, 1st edition. ed. Cambridge University Press, Cambridge.

Kay, T., Keller, L., Lehmann, L., 2020. The evolution of altruism and the serial rediscovery of the role of relatedness. Proc. Natl. Acad. Sci. 117, 28894–28898. 10.1073/pnas.2013596117

Kello, C.T., Bella, S.D., Médé, B., Balasubramaniam, R., 2017. Hierarchical temporal structure in music, speech and animal vocalizations: jazz is like a conversation, humpbacks sing like hermit thrushes. J. R. Soc. Interface 14, 20170231. 10.1098/rsif.2017.0231

Kilner, J.M., Lemon, R.N., 2013. What We Know Currently about Mirror Neurons. Curr. Biol. 23, R1057–R1062. 10.1016/j.cub.2013.10.051

Melis, A.P., 2018. The evolutionary roots of prosociality: the case of instrumental helping. Curr. Opin. Psychol., Early Development of prosocial behavior 20, 82–86. 10.1016/j.copsyc.2017.08.019

Merker, B., Morley, I., Zuidema, W., 2015. Five fundamental constraints on theories of the origins of music. Philos. Trans. R. Soc. B Biol. Sci. 370, 20140095. 10.1098/rstb.2014.0095

Mohd Nor Ihsan, N.S., Abdul Sani, S.F., Looi, L.M., Cheah, P.L., Chiew, S.F., Pathmanathan, D., Bradley, D.A., 2023. A review: Exploring the metabolic and structural characterisation of beta pleated amyloid fibril in human tissue using Raman spectrometry and SAXS. Prog. Biophys. Mol. Biol. 182, 59–74. 10.1016/j.pbiomolbio.2023.06.002

Peretz, I., Vuvan, D., Lagrois, M.-É., Armony, J.L., 2015. Neural overlap in processing music and speech. Philos. Trans. R. Soc. B Biol. Sci. 370, 20140090. 10.1098/rstb.2014.0090

Pfattheicher, S., Nielsen, Y.A., Thielmann, I., 2022. Prosocial behavior and altruism: A review of concepts and definitions. Curr. Opin. Psychol. 44, 124–129. 10.1016/j.copsyc.2021.08.021

Raimondi, T., Di Panfilo, G., Pasquali, M., Zarantonello, M., Favaro, L., Savini, T., Gamba, M., Ravignani, A., 2023. Isochrony and rhythmic interaction in ape duetting. Proc. Biol. Sci. 290, 20222244. 10.1098/rspb.2022.2244

Ravignani, A., Bowling, D.L., Fitch, W.T., 2014. Chorusing, synchrony, and the evolutionary functions of rhythm. Front. Psychol. 5, 1118. 10.3389/fpsyg.2014.01118

Rothenberg, D., Roeske, T.C., Voss, H.U., Naguib, M., Tchernichovski, O., 2014. Investigation of musicality in birdsong. Hear. Res. 308, 71–83. 10.1016/j.heares.2013.08.016

Sacks, O., 2008. Musicophilia: Tales of Music and the Brain, Revised and Expanded Edition, Revised&enlarged edition. ed. Vintage, New York.

Silk, J.B., House, B.R., 2011. Evolutionary Foundations of Human Prosocial Sentiments, in: In the Light of Evolution: Volume V: Cooperation and Conflict. National Academies Press (US).

Snowdon, C.T., Teie, D., 2009. Affective responses in tamarins elicited by species-specific music. Biol. Lett. 6, 30–32. 10.1098/rsbl.2009.0593

Sornette, D., 2006. Critical Phenomena in Natural Sciences: Chaos, Fractals, Selforganization and Disorder: Concepts and Tools, 2nd edition. ed. Springer, Berlin, Heidelberg.

Spitzer, M., 2021. The Musical Human: A History of Life on Earth. Bloomsbury Publishing USA, New York.

Takahashi, R., Miller, J.H., 2007. Conversion of amino-acid sequence in proteins to classical music: search for auditory patterns. Genome Biol. 8, 405. 10.1186/gb-2007-8-5-405

Tay, N.W., Liu, F., Wang, C., Zhang, H., Zhang, P., Chen, Y.Z., 2021. Protein music of enhanced musicality by music style guided exploration of diverse amino acid properties. Heliyon 7. 10.1016/j.heliyon.2021.e07933

Trainor, L.J., 2015. The origins of music in auditory scene analysis and the roles of evolution and culture in musical creation. Philos. Trans. R. Soc. B Biol. Sci. 370, 20140089. 10.1098/rstb.2014.0089

Trehub, S.E., Becker, J., Morley, I., 2015. Cross-cultural perspectives on music and musicality. Philos. Trans. R. Soc. B Biol. Sci. 370, 20140096. 10.1098/rstb.2014.0096

Trimble, M., Hesdorffer, D., 2017. Music and the brain: the neuroscience of music and musical appreciation. BJPsych Int. 14, 28–31.

Trost, W., Trevor, C., Fernandez, N., Steiner, F., Frühholz, S., n.d. Live music stimulates the affective brain and emotionally entrains listeners in real time. Proc. Natl. Acad. Sci. U. S. A. 121, e2316306121. 10.1073/pnas.2316306121

Wang, S., 2019. Efficiently Calculating Anharmonic Frequencies of Molecular Vibration by Molecular Dynamics Trajectory Analysis. ACS Omega 4, 9271–9283. 10.1021/acsomega.8b03364

When Bjork Met Attenborough, 2014.. Sony.

Wigner, E.P., 1960. The unreasonable effectiveness of mathematics in the natural sciences. Richard courant lecture in mathematical sciences delivered at New York University, May 11, 1959. Commun. Pure Appl. Math. 13, 1–14. 10.1002/cpa.3160130102

Wilson, D.S., Wilson, E.O., 2007. Rethinking the theoretical foundation of sociobiology. Q. Rev. Biol. 82, 327–348. 10.1086/522809

Wilson, E.O., Hölldobler, B., 2005. Eusociality: Origin and consequences. Proc. Natl. Acad. Sci. U. S. A. 102, 13367–13371. 10.1073/pnas.0505858102

Wilson, E.O., Nowak, M.A., 2014. Natural selection drives the evolution of ant life cycles. Proc. Natl. Acad. Sci. U. S. A. 111, 12585–12590. 10.1073/pnas.1405550111

Wolfe, J., 2002. Speech and Music, Acoustics and Coding, and What Music Might Be “For.”

Yu, C.-H., Qin, Z., Martin-Martinez, F.J., Buehler, M.J., 2019. A Self-Consistent Sonification Method to Translate Amino Acid Sequences into Musical Compositions and Application in Protein Design Using Artificial Intelligence. ACS Nano 13, 7471–7482. 10.1021/acsnano.9b02180

