## Supplemental Methods for "Musicality in protein interaction dynamics informs the multi-scale evolution of prosocial behavior"

#### PDB structure and model preparation

Protein structures of amyloid fibrils and protein-ligand interactions (Table 1) were obtained from the Protein Data Bank (PDB). After downloading the structures from the PDB database, any crystallographic reflections, ions, and other solvents used in the crystallization process were removed. Any missing loop structures in the protein structures were inferred using the protocol for loop refinements on the ModLoop webserver at The Anderj Sali lab at UCSF. The pdb4amber (AmberTools20) was also employed to add hydrogen atoms (i.e. reduce the structure) and remove crystallographic waters (Case et al., 2005). All molecular color-mapping of our results were conducted in UCSF ChimeraX (Goddard et al., 2018; Pettersen et al., 2021). To prepare the molecular dynamics comparisons for ATOMDANCE (Babbitt et al., 2024), we created unbound protein structures by removing the small molecule ligands, protein partners or in the case of amyloids we removed every other adjacent fibril creating isolated protein strands surrounded by the solvent in the system. Thus an amyloid complex with 12 fibrils would be prepped into a system with 6 separate spaced fibrils and compared to a system with 6 compacted fibrils.

#### Molecular dynamic simulation protocols

For each molecular dynamic comparison (i.e. amyloid fibrils bound vs. unbound to each other or proteins bound vs. unbound to small molecule ligands) simulations were performed. In brief, for each MD comparison, large replicate sets of accelerated MD simulation were prepared and then conducted using the particle mesh Ewald method (Darden et al., 1993; Ewald, 1921) implemented on NVIDIA graphical processor units under Langevin integration using mixed precision implementation via OpenMM (Eastman et al., 2023, 2013). The MD simulations were done on a high performance computing workstation mounting dual Nvidia 2080Ti graphics processor units. All comparative MD analysis via our ATOMDANCE was based upon 100 randomly resampled windows collected on of 10ns of accelerated MD in each comparative state (i.e. bound vs. unbound). Explicitly solvated protein systems were first prepared using teLeap (AmberTools 20) (Case et al., 2005), using ff14SSB protein force field (Maier et al., 2015), in conjunction with other forcefields as needed including DNA.OL15, and GAFF2 small molecule force field (Wang et al., 2004) modified via the sqm program in antechamber (AmberTools 20)(Wang et al., 2006). Solvation was generated using the Tip3p water model in a 12nm octahedral water box. Charge neutralization was performed using Na+ and Cl- ions using the AmberTools22 teLeap program. The sqm program (version 17) in antechamber software (AmberTools 20) was used to perform semiempirical quantum mechanical optimization to obtain the force field modifications needed to run molecular dynamics simulation on small molecular ligand interactions with proteins. For each MD comparison, an energy minimization was first performed, then heated to 300K for 300 pico seconds, followed by 10 ns of equilibration, and then finally a replicate set of 100 MD production run was created for each comparative state. Each MD production run was simulation for 1 ns of time and sampled at 5000 frames. All simulations were regulated using the Andersen thermostat at 300k and 1atm (Andersen, 1980). Root mean square atom fluctuations were conducted in CPPTRAJ using the atomicfluct command (Roe and Cheatham, 2013).

Table 1. PDB IDs and descriptions for the amyloid fibrils and protein-ligand interactions used in the molecular dynamic simulations

| PDB | CATEGORY | DESCRIPTION |
|---|---|---|
| 2a3w | amyloid fibril | Human serum amyloid p-component |
| 2lbu | amyloid fibril | HET-s amyloid |
| 2mus | amyloid fibril | HET-s amyloid |
| 6dso | amyloid fibril | AA amyloid (murine) |
| 6ic3 | amyloid fibril | AL amyloid (lambda 1light chain) |
| 6lni | amyloid fibril | Human prion amyloid |
| 6uur | amyloid fibril | Human prion amyloid variant |
| 6zcf | amyloid fibril | SAA1.1 amyloid (murine) |
| 6zrf | amyloid fibril | IAPP-islet amyloid (amylin) |
| 6zrr | amyloid fibril | IAPP-islet amyloid (amylin) |
| 7qv6 | amyloid fibril | Aurein 3.3 amyloid |
| 7rl4 | amyloid fibril | PrP23-144 amyloid (human) |
| 7td6 | amyloid fibril | aRML prion (murine) |
| 7umq | amyloid fibril | Type 1 prion (human GSS disease) |
| 7yat | amyloid fibril | Hamster prion 108-144 |
| 7zir | amyloid fibril | hnRNPDL amyloid |
| 7zky | amyloid fibril | AA amyloid (human) |
| 8a00 | amyloid fibril | ME7 scrapie prion (murine) |
| 8az4 | amyloid fibril | IAPP-islet amyloid 2PF-L (human) |
| 8dja | amyloid fibril | PrP23-144 amyloid (murine) |
| 8efu | amyloid fibril | A22L prion (murine) |
| 8enq | amyloid fibril | CsgA fibril (E. coli) |
| 8g2v | amyloid fibril | LECT2 amyloid (human) |
| 8r4a | amyloid fibril | Lysozyme variant fibril (human) |
| 8spa | amyloid fibril | CPEB3 prion (human) |
| 8wzx | amyloid fibril | Hamster prion 23-144 |
| 9dmy | amyloid fibril | Chronic wasting disease prion |
| 1j6z | protein-ligand | uncomplexed actin |
| 1pkg | protein-ligand | c-Kit kinase |
| 1qz5 | protein-ligand | Nitril hydratase (pseudomonas) |
| 1uwh | protein-ligand | B-Raf kinase (human) |
| 1uwj | protein-ligand | B-Raf kinase variant (human) |
| 3eeb | protein-ligand | RTX cysteine protease (Vibrio cholera) |
| 4fal | protein-ligand | 17beta-hydroxysteroid dehydrogenase |
| 1cdw | protein-DNA | TATA binding protein (human) |
| 1qna | protein-DNA | TATA binding protein (Arabidopsis) |
| 1blb | protein-protein | Eye lens beta-crystallin complex |
| 1fgb | protein-protein | Enterotoxin complex (Vibrio cholera) |
| 1kx5 (2 chain) | protein-protein | Nucleosome core complex (human) |

| | | |
|---|---|---|
| **1kx5 (4 chain)** | **protein-protein** | **Nucleosome core complex (human)** |
| **1ubq** | **protein-protein** | **uncomplexed ubiquitin** |
| **5vix** | **protein-protein** | **ubiquitin dimer** |
| **6m17** | **protein-protein** | **SARS-CoV-2 RDB and ACE-2 (human)** |
| **6nxl** | **protein-protein** | **Ubiquitin binding variants** |

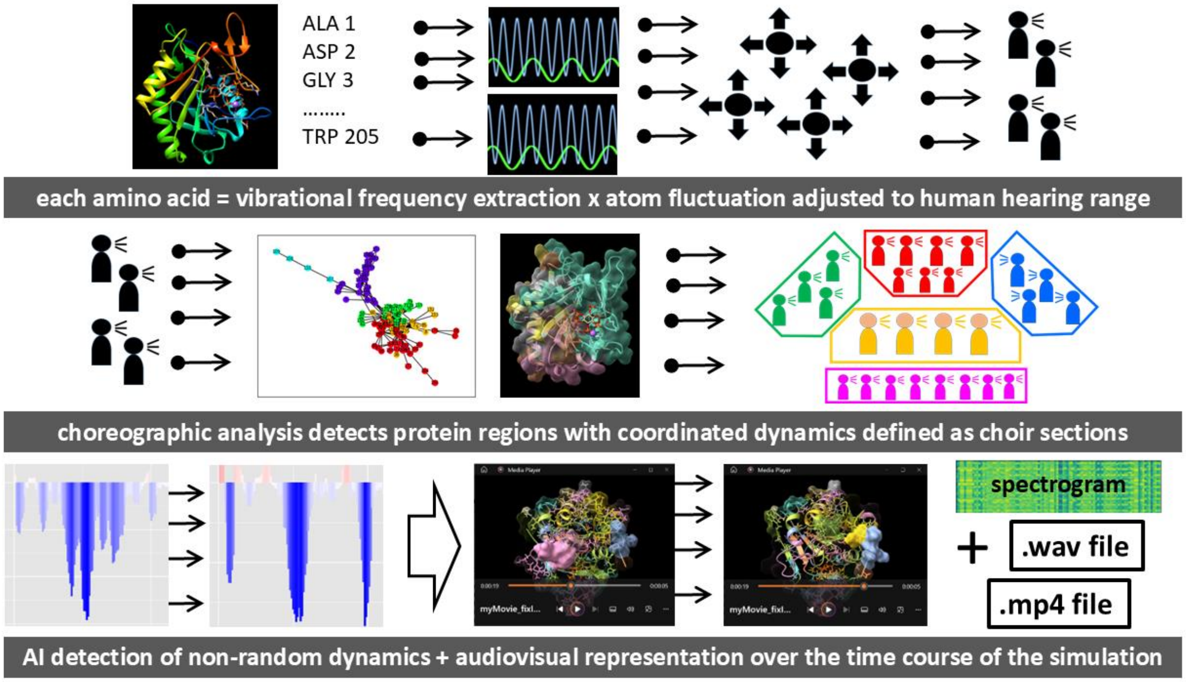


**Figure 2A. Graphical overview of the data sonification of protein interaction dynamics.** Initially, vibrational frequencies are extracted per each amino acid and adjusted in accordance to average atom fluctuation and the pitch range of human hearing. Using choreographic analysis in ATOMDANCE, each amino acid frequency (i.e. individual voice) is layered in accordance to a community with which it shares coordinated dynamics (i.e. choir section). Using maxDemon denoising in ATOMDANCE, each frequency volume is adjusted according to its level of non-random shift in motion (i.e. functional binding) during protein interaction. Sound files and movie files are created by running this overall analysis pipeline over sub-segments of the molecular dynamic trajectory, subsequently creating each individual movie frame.

Example here – BRAF kinase domain interaction with the cancer drug sorafenib

https://people.rit.edu/gabsbi/img/videos/AAVexample.mp4

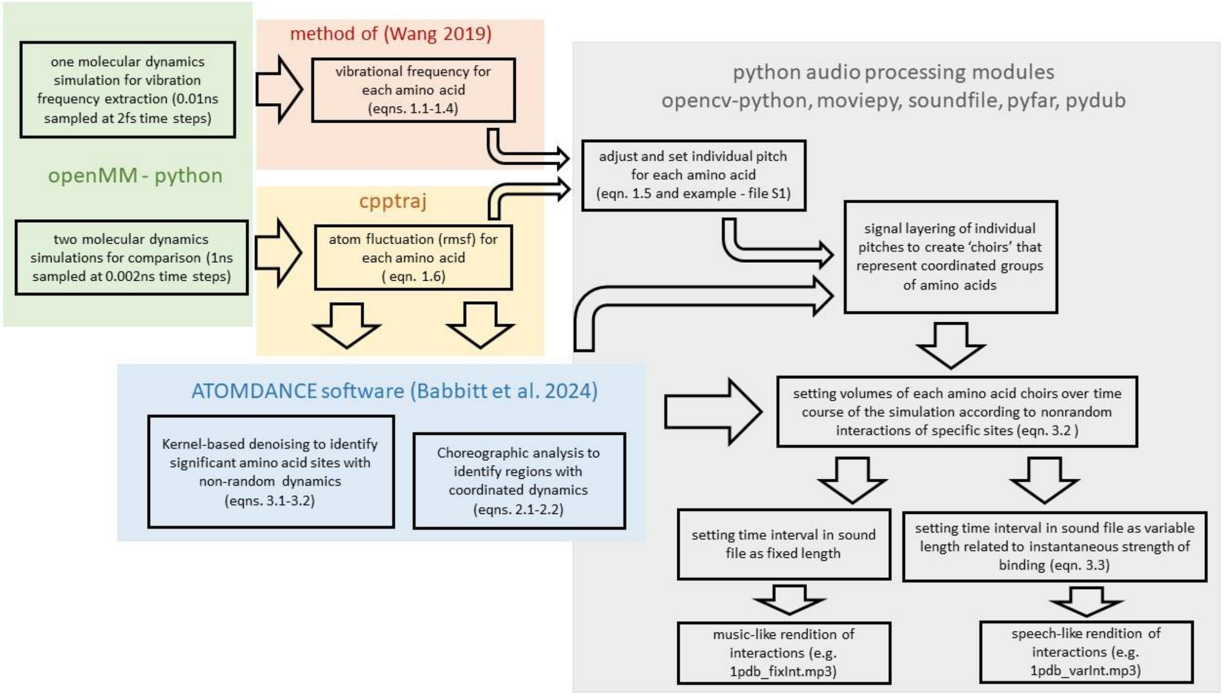


**Figure 2B. Schematic overview of the data sonification of protein interaction dynamics.** All the methods and equations are fully defined in the Supplemental Methods section.

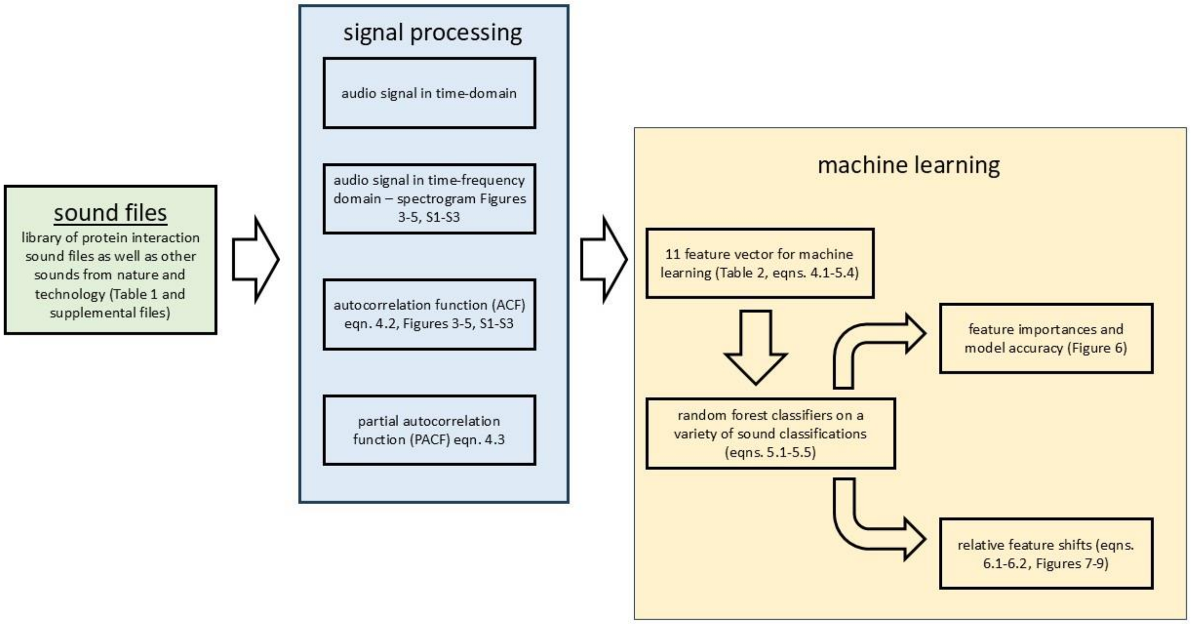


**Figure 2C. Schematic overview of the machine learning analysis of the sounds of protein interaction dynamics in the context of natural sounds.** All the methods and equations are fully defined in the Supplemental Methods section.

#### Vibrational frequency extraction and pitch adjustment of each amino acid to human hearing range

To keep the AAV pipeline computationally efficient, the vibrational frequencies are extracted from a relatively short but frequently sampled MD production run (0.01ns time stepped every 2fs and sampled every 2fs) taken after a much longer but less frequently sampled period of equilibration (typically 10-100 ns time stepped every 2fs and sampled every 1ps). Following the method of (Wang 2019), the Eckart frame processes molecular dynamics (MD) data to align it in a manner that isolates internal motions from external motions, such as overall translations and rotations. The input data is a three-dimensional array with dimensions, where is the number $(n, m, 3)$ $n$ of frames and $m$ is the number of atoms per frame. In summary, the original trajectory is transformed via a spectral decomposition of the total covariance matrix across all frames, from which the orthogonal rotation matrix R is obtained. Thus the total variation across frames is

$$C_t = R\,\Lambda\,R^T \qquad 1.1$$

Singular value decomposition (SVD) is then performed on the total covariance matrix to find the rotation matrix that minimizes the differences between the frames. This technique decomposes the total covariance matrix into its principal components, which allows for the identification of the primary modes of internal motion in the MD data.

$$U, S, V^T = SVD(C_t) \qquad 1.2$$

$U$ and $V^T$ are unitary matrices, i.e., square matrices of complex numbers, and S is a diagonal matrix containing the singular values. The Eckart frame rotation matrix is given by the first $m$ columns of $V^T$ as $R = V^T$. The rotation matrix is then applied to each frame to align all frames to a common reference, which isolates internal atomic motions.

$$x''_{ij} = x_{ij}R^T \qquad 1.3$$

Where $x_{ij}$ is the spatial position of the jth atom of frame i. The output is the Eckart-rotated MD data with the same shape as the input, effectively aligning the frames. Subsequently, the Fast Fourier Transform (FFT) is then applied to the rotated data to convert it from the time domain to the frequency domain. Note that the rotated data and sampling rate $f_s$ are the inputs for this technique. The FFT is then performed on each frame of the rotated data along the axis of the atoms.

$$F_i = FFT(x''_i)f_s \qquad 1.4$$

The FFT transforms the time-domain data into the frequency domain, featuring the dominant frequencies and periodicities in the motion of the atoms. This workflow enables extraction of rapid molecular dynamics in the femtosecond scale range by separating the intrinsic motions of the system within each amino acid without the influence of longer range translations and rotations due to larger scale motions of the entire protein in the solvent.

After the extraction of the vibrational frequency, a .wav file is created for each amino acid signal. To adjust the original signal to the variable range of pitch best detected by human hearing, while still retaining information about the specific molecular dynamics of a given amino acid, the original signal files were raised by up to a maximum of 13 octaves (O) in relation to *rmsf* for each amino acid

$$O = 9 + 20 \cdot (\overline{rmsf}) \qquad 1.5$$

where mean rmsf is atom fluctuation averaged over the 4 backbone atoms in the amino acid is given by

$$\overline{rmsf} = \frac{1}{m}\sum_{i=1}^{m}\sqrt{\left(\frac{1}{n} * \sum_{j=1}^{n}\left(\left(s_{ijx} - w_x\right)^2 + \left(s_{ijy} - w_y\right)^2 + \left(s_{ijz} - w_z\right)^2\right)\right)} \qquad 1.6$$

where $n$ is the number of frames in the MD simulation and $m$ is the number of amino acid backbone atoms per frame (i.e. $m = 4$) and where $s$ represents the set of XYZ atom coordinates for the $m$ atoms for a given amino acid residue over the $n$ time points, and $w$ represents the center of the overall coordinate dimensional structure during the MD production run for a given amino acid. This adjustment to the pitch of the original vibrational frequencies extracted by eqns. 1.1-1.7 creates a range of unevenly distributed pure tones for amino acids that is reflective of the overall magnitude of the regional dynamic motion in different parts of the protein. Note that these tones are reflective of local dynamics and not the identity of the amino acid itself.

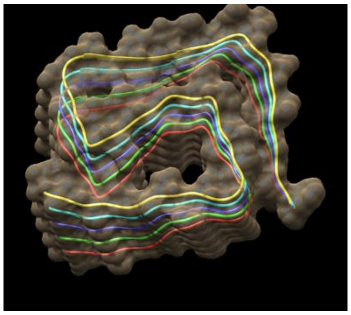

PDB:8spa B amyloid structure.

Examples of these adjusted pitches for each of the amino acids present in the amyloid PDB:8spa shown above are given in supplementalData (DOI: 10.5281/zenodo.15013666) in the exampleSignalAdjust subfolder.

**Choreographic analysis to identify regions with coordinated dynamics and establish layering of sounds of the choir sections**

In the MD simulation, every site μ on the protein is compared to every site *v* using a mixed effects model ANOVA where atom fluctuation represents a fixed effect (α) in the model and a time sample represents a random effect (β) in the model (with mean(rmsf) = mean fluctuation within a given time interval and ε= error or residual term). Thus the general linear model becomes

$$f(\overline{rmsf})_{ct} = \overline{rmsf} + \alpha_c + \beta_t + \alpha\beta_{ct} + \varepsilon_{ct} \qquad 2.1$$

where c represents the site class ($\mu$ or ν) and $t$ represents the random time sampling group collected by the cpptraj program.

For the choreographic analysis, the p-value of interaction between atom fluctuation levels between site μ and site *v* and the time subsamples in the MD simulation (i.e. $\alpha\beta_{ct}$) indicates the significance of an interaction of fixed differences in atom fluctuation between the two separate sites over time (i.e. a coordinated or choreographed physical motion). The p-values are corrected for false discovery rate via Benjamini Hochberg method and shown as a heat map representing significance of areas of synchronously shifting site dynamics on the protein. In the second step of the analysis, intended to define communities of coordinated regions of protein dynamics, the strongest interaction p-values for all site μ to site v comparisons are represented by a graph network where ($k_n$), the degree of node n is

$$k_n = A_{\mu\nu} \sum (p_\mu p_\nu)_i \qquad 2.2$$

where $p_\mu$ and $p_\nu$ are the interaction p values for sites $\mu$ and $\nu$, and $A_{\mu\nu}$ is the adjacency matrix connecting nodes $p_\mu$ and $p_\nu$ . An autotuned cutoff that collects the 5% strongest interaction p-values was used. The Louvain community detection algorithm iterates a two-step process of modularity optimization followed by community aggregation until community identities of all nodes are stable. It is implemented in our code by the python package NetworkX ("Proceedings of the Python in Science Conference (SciPy): Exploring Network Structure, Dynamics, and Function using NetworkX," n.d.). In accordance with each community detected, the sounds of the amino acids in that community were looped and overlayed using 0 positional shift and a gain during overlay of -4. A Butterworth bandpass filter was applied over the range of 500-15000Hz.

**Sound sequence generation from extraction of non-random binding dynamics inferred from the comparison of bound vs. unbound states**

During the course of each MD production run, 100 intervals were individually processed and analyzed as individual movie frames. Withing each frame a comparative analysis of the local MD simulation was conducted for isolating single sites responsible for non-random dynamics during binding interactions. A Gaussian process kernel learner was applied to isolate maximum mean discrepancy (MMD) between learned features of atom fluctuation that shift between bound and unbound states of the protein, identifying key sites involved in functional binding. This analysis uses site-wise training of Gaussian processes machine learners with tuned radial basis kernel functions to specify MMD in reproducing kernel Hilbert space (RKHS) that describes the distance in learned features between the two protein dynamic states at all given sites on the protein. Thus, the kernel function describing the mapping of the local *rmsf* values ($u_i$ and $u_j$) being compared across the protein's bound vs. unbound functional state is

$$k(u_i, u_j) = exp\left(\frac{\|u_i - u_j\|^2}{2\sigma^2}\right) \qquad 3.1$$

Note that σ is sampled derived. And the empirical estimations of MMD, or distance between feature means is given by

$$MMD^2(U,V) = \frac{1}{m(m-1)}\sum_i \sum_{j\neq i} k(u_i, u_j) - 2\frac{1}{m(m-1)}\sum_i \sum_j k(u_i, v_j) + \frac{1}{m(m-1)}\sum_i \sum_{j\neq i} k(v_i, v_j) \qquad 3.2$$

where u's are the original local *rmsf* values and v's are generated values evaluated on the kernel and m is 4 (the number of atoms in the amino acid backbone).

The learners are trained using a local atom fluctuation feature vector comprised of *rmsf* values from sites -2, -1, 0, 1, 2 positions on the protein chain relative to the site being analyzed. A key concept here is that because the learner cannot optimize on random differences in atom fluctuation caused by thermal noise it acts as a noise filter, thus eliminating motion dampening that is not directly due to non-random differences in atom fluctuation between the sites being compared (i.e. functional aspects of molecular interactions directly involved in the binding interaction)

The volume of the voice of these key sites driving the functional dynamics (i.e. nonrandom dynamics), along with each voice in their own choir section were adjusted according to the level of MMD at the given site. Thus, key sites will both activate and dominate the voice of the choir section to which they belong. In the resulting movies, the relative level of MMD is represented by the opacity of the surface shown for a given amino acid. Sidechains are shown whenever the transparency is less than 80%. A wav file with an overlay of the active choirs is generated.

Lastly, the AAV pipeline produces two types of renditions of sound over time. One of these is more musical in that it has fixed intervals of 250 milliseconds for each frame in the analysis over the time course of the molecular dynamics simulation, while the other is potentially more speech-like in that time intervals in the sound sequence are an inverse function of the strength of binding at the specific time points in the simulation. Thus the interval (in milliseconds) for each frame in the time course of the molecular dynamics simulation is

$$interval = 250 - 250 * \sum_{i=1}^{L} MMD_i \qquad 3.3$$

Graphical and schematic overviews of the data sonification procedure in AAV is shown in Figure 2A and 2B.

**Signal and spectral correlation analyses of the sound files to define acoustic features**

Resulting sound files for the protein interactions were post-processed using spectral correlation analysis of the spectrogram representing variation of sound in the time and frequency domains. To extract the acoustic complexity (i.e. note variability) following the method of Sawant et al. 2021 for analyzing bird song complexity, the spectral cross-correlation is converted to a Note Variability Index (NVI) ranging between 0 and 1 where

$$Note\ Variability\ Index\ (NVI) = variability = \frac{\sum_{i=1}^{N_C}\sum_{j=1}^{N_C}\left(1 - c_{\Delta T_{I,j}}\right)}{N_C(N_C - 1)} \qquad 4.1$$

forming an $N_C$ X $N_C$ similarity matrix where $N_C$ is the number of notes and $C_{\Delta T}$ is the association between two notes. Thus, a lack of cross-correlation across all pairs of notes or data columns in a spectrogram represents high variability in the sound file while a tone repeated constantly in the time domain has high cross-correlation but no complexity. However, it is important to understand that while variability in the signal increases to the point of unpredictability, the variability eventually converts from a complex pattern to memoryless white noise (where the next value in a sequence cannot be predicted at all).

The autocorrelation function (ACF) with time lag (k) is also computed across the time domain (x) as

$$ACF = p(k) = \frac{cov[x_t, x_{t-k}]}{cov[x_t, x_t]} \qquad 4.2$$

A partial autocorrelation function (PACF) is also generated as well.

$$PACF = p(k) = \frac{cov[x_t, x_{t-k-b}]}{cov[x_t, x_t]} \qquad 4.3$$

Where b is the best linear predictor of $x_t$ based upon the previous k-1 values of the lag term k.

Several measures of acoustic periodicity are developed. A peak detection algorithm is applied to the ACF using a height and width of (6,8) to collect the distinct number of peaks (N) in the ACF. As this count ranges from the single digits to often tens of thousands we log transform the count and defined it as the acoustic signal harmonic layering.

$$number\ of\ autocorrelation\ peaks = level\ of\ signal\ layering = \left(N_{ACF}^{peaks}\right) \quad 4.4$$

This simple measure of the peak counts in the ACF also strongly correlates to another indicator of periodicity taken as the level of evenness in the time lags between peaks (L) taken over all significant peaks in the ACF calculated as

$$evenness\ index = periodicity = \sum_{i=1}^{n} {(L_i - \bar{L})^2}/{n} \quad 4.5$$

Additionally, the r value at the first order of the ACF function is taken as an indicator of echo or reverberation, and the order of the moving average (MA order) is also taken as an indicator of distortion due to either layering or noise. While the lag value at the first peak varies in time depending upon the signal being analyzed, it represents the most immediate and shortest repetitive acoustic element in the signal. Strong short-term memory implies that at lag period = 1, the current values of the frequency domain in the spectrogram are strongly predicted by the prior values of the frequency domain at the most immediate lag value. For example, a highly repetitive stationary signal will have high 1st order memory as will Brown noise, where even though the time-series progression is random, it is completely depending upon the immediately preceding value in the random walk. From an acoustic perspective, strong reverberation (short-term memory) implies that a sound is highly non-random in its transitions over its shortest meaningful time period.

$$first\ order\ autocorrelation = reverberation = r_{L\,=\,1} \quad 4.5$$

The longest lag distance to the on the ACF function where the AC is still significant is known as the order of the moving average (MA) is also used as a feature as well

$$order\ of\ moving\ average\ (MA) = lengthspacing\ of\ signal\ layering = \mathrm{maximum}(L_{acf}) \quad 4.6$$

Several measures from the partial autocorrelation function (PACF) are used as well. These are the order of the autoregression (AR) and the autocorrelation level observed at the order of the AR taking as a measure of acoustic signal delay and level of repetitiveness, respectively.

$$order\ of\ autoregression\ (AR) = length\ spacing\ of\ signal\ delay = \mathrm{maximum}(L_{pacf}) \quad 4.7$$

$$AC\ at\ order\ of\ autoregression = level\ of\ signal\ delay = (r_{pacf})_{L\,=\,max} \quad 4.8$$

We also measure the long-term memory or persistence of the acoustic signal as

$$memory\ level\ index = persistence = 2\big(abs(0.5 - H)\big) \quad 4.9$$

where H is the Hurst exponent whose values are interpreted as 0<H<0.5 is a mean-reverting or trend-resistant signal, and 0.5>H>1 is a predictably evolving or trending signal, and H = 0.5 is a random signal with no long-term memory. Our memory index conversion of the Hurst exponent scales the long-term memory from 0 to 1 (non-existent to strong) while disregarding whether it is mean reverting vs. trending. From an acoustic perspective, strong long-term memory implies that sounds are persistently evolving (or failing to evolve) over many time periods in a complex way. Lastly, and perhaps most importantly, we include the test statistics from the augmented Dickey-Fuller test as a measure of non-stationarity of the acoustic signal. The ADF statistic grows negatively from zero as the information content increases in a signal, thus it is often used to measure the level of information present in speech In music it indicates fluctuation of volume or pitch. The PACF plots assume stationarity of signal and remove signal layering as one moves outwards on the ACF plots. Thus, in an acoustic context, autoregression based upon PACF captures more of the delay in the signal, whereas the ACF captures the overall layering of the acoustic signal.

A summary of the 11 acoustic features and their interpretations is given in Table 2.

**Machine learning classification of music and other natural sounds**

These features are used to train subsequent random forest models that were used to assess comparative classification of sound types derived from our protein sonification when compared within the context of human speech, music, and other natural and computer-generated sounds. All sound files used are listed in featureData_allFolders.dat and folder_list.txt totaling 1185 sound samples (included in SupplementalData (DOI: 10.5281/zenodo.15013666). Note that some music files are omitted due to copyright, but the titles of the music tracks analyzed can be seen in the first column of featureData_allFolders.dat. Our random forests consisted of 500 decision trees using a 80/20% training/testing data split. It is implemented with python scikit-learn RandomForestClassifier function which implements the ID3 with CART algorithm to create each decision tree. Splits on the trees are called using Gini impurity or Gini index.

$$Gini\ impurity = \sum_{i-1}^{N} f_i(1-f_i) \qquad 5.1$$

Where $f_i$ is the frequency of label i at a node and N is the number of labels.

The importance for each feature (I) in the random forest is given by

$$I = \frac{\sum_{j=all\ trees} norm\ I_j}{T} \qquad 5.2$$

where the normalized feature importance is for a give tree is

$$norm\ I_{ij} = \frac{I_i}{\sum_{j=all\ features} I_j} \qquad 5.3$$

where the feature importance for a give tree is

$$I_i = \frac{\sum_{j:node\ j\ splits\ on\ feature\ i} ni_j}{\sum_{k=all\ nodes} ni_k} \qquad 5.4$$

where $n_j$ the importance of each node j is

$$n_j = w_j G_j - w_{j(left)} G_{j(left)} - w_{j(right)} G_{j(right)} \qquad 5.5$$

where w is the number of samples reaching node j, G is the impurity value of node j and left(j) and right(j) indicate the child nodes of node j

**Table 2. Feature vector supplied to the random forest classifiers used to discern protein generated from musical and non-musical sounds. Each acoustic feature is described in its potential contribution or relation to musicality.**

| ACOUSTIC FEATURE | MEASURES/DETECTS | RELATION TO MUSICALITY | DERIVATION |
|---|---|---|---|
| dominant frequency | most common pitch | voice and instrument pitch | time domain |
| Note Variability Index (NVI) | variability of spectrogram | melodic and harmonic complexity = predictability, surprise, information | frequency and time domains |
| ADF statistic | stationarity | fluctuation of volume or pitch | time domain |
| Hurst exponent | mean-reverting vs. trending | evolution, progression, development | time domain |
| Memory Level Index (MLI) | randomness vs. persistence | evolution, progression, development | time domain |
| first-order autocorrelation | echo or reverberation | dimension, ambience, distortion | ACF |
| order moving average (MA) | length of signal layering | harmonic structure and progression | ACF |
| log N autocorrelation peaks | level of signal layering | harmonic structure and progression | ACF |
| Evenness Index | signal periodicity | rhythmic structure and cadence | ACF |
| order autoregression (AR) | length of signal delay | rhythmic structure and cadence | PACF |
| correlation (order of AR) | level of signal delay | rhythmic structure and cadence | PACF |

**Determination of relative acoustic feature shift via Jenson-Shannon distance**

We measure the relative acoustic feature shifts (ΔF) of pairs of the normalized acoustic feature values (e.g. male vs female speech) towards or away from a third reference feature (e.g. music or animal vocalizations) by calculating the respective differences in the Jensen-Shannon Distance ($D_{js}$) from the reference feature.

$$feature\ shift = \Delta F = D_{js}(P_1||Q) - D_{js}(P_2||Q) \qquad 6.1$$

where P1 and P2 are the features to be compared relative to the values of feature Q and

$$D_{js}(P||Q) = \frac{1}{2}D_{kl}(P||M) + \frac{1}{2}D_{kl}(Q||M) \qquad 6.2$$

where M = 0.5(P+Q) is a mixture distribution of P and Q and $D_{kl}$ is the Kullback-Leibler divergence of P from Q or (P||Q). Empirical p-values for each feature shift were determined using a permutation test (n = 100000) on the bootstrapped data.

A schematic overview of the analysis of the sound files is given in Figure 2C.

The AAV pipeline code is found in the ATOMDANCE code repo at

https://github.com/gbabbitt/ATOMDANCE-comparative-protein-dynamics

with website

https://gbabbitt.github.io/ATOMDANCE-comparative-protein-dynamics/

All post processing codes used in the analyses are in the postProcessingCode folder in SupplementaryData. All plots generated during post-processing are also present in supplementalData (subfolders are acousticFeatureExtraction, exampleSignalAdjust, featureShiftAnalysis, postProcessingCode, and RFclassification&comparison). This data is reposited at Zenodo.org under digital object identifier (DOI: 10.5281/zenodo.15013666).

### References

Andersen, H.C., 1980. Molecular dynamics simulations at constant pressure and/or temperature. J. Chem. Phys. 72, 2384–2393. https://doi.org/10.1063/1.439486

Babbitt, G.A., Rajendran, M., Lynch, M.L., Asare-Bediako, R., Mouli, L.T., Ryan, C.J., Srivastava, H., Rynkiewicz, P., Phadke, K., Reed, M.L., Moore, N., Ferran, M.C., Fokoue, E.P., 2024. ATOMDANCE: Kernel-based denoising and choreographic analysis for protein dynamic comparison. Biophys. J. 123, 2705–2715. https://doi.org/10.1016/j.bpj.2024.03.024

Blondel, V.D., Guillaume, J.-L., Lambiotte, R., Lefebvre, E., 2008. Fast unfolding of communities in large networks. J. Stat. Mech. Theory Exp. 2008, P10008. https://doi.org/10.1088/1742-5468/2008/10/P10008

Case, D.A., Cheatham, T.E., Darden, T., Gohlke, H., Luo, R., Merz, K.M., Onufriev, A., Simmerling, C., Wang, B., Woods, R.J., 2005. The Amber biomolecular simulation programs. J. Comput. Chem. 26, 1668–1688. https://doi.org/10.1002/jcc.20290

Darden, T., York, D., Pedersen, L., 1993. Particle mesh Ewald: An N·log(N) method for Ewald sums in large systems. J. Chem. Phys. 98, 10089–10092. https://doi.org/10.1063/1.464397

Eastman, P., Friedrichs, M.S., Chodera, J.D., Radmer, R.J., Bruns, C.M., Ku, J.P., Beauchamp, K.A., Lane, T.J., Wang, L.-P., Shukla, D., Tye, T., Houston, M., Stich, T., Klein, C., Shirts, M.R., Pande, V.S., 2013. OpenMM 4: A Reusable, Extensible, Hardware Independent Library for High Performance Molecular Simulation. J. Chem. Theory Comput. 9, 461–469. https://doi.org/10.1021/ct300857j

Eastman, P., Galvelis, R., Peláez, R.P., Abreu, C.R.A., Farr, S.E., Gallicchio, E., Gorenko, A., Henry, M.M., Hu, F., Huang, J., Krämer, A., Michel, J., Mitchell, J.A., Pande, V.S., Rodrigues, J.P., Rodriguez-Guerra, J., Simmonett, A.C., Singh, S., Swails, J., Turner, P., Wang, Y., Zhang, I., Chodera, J.D., Fabritiis, G.D., Markland, T.E., 2023. OpenMM 8: Molecular Dynamics Simulation with Machine Learning Potentials. ArXiv arXiv:2310.03121v2.

Ewald, P.P., 1921. Die Berechnung optischer und elektrostatischer Gitterpotentiale. Ann. Phys. 369, 253–287. https://doi.org/10.1002/andp.19213690304

Goddard, T.D., Huang, C.C., Meng, E.C., Pettersen, E.F., Couch, G.S., Morris, J.H., Ferrin, T.E., 2018. UCSF ChimeraX: Meeting modern challenges in visualization and analysis. Protein Sci. Publ. Protein Soc. 27, 14–25. https://doi.org/10.1002/pro.3235

Maier, J.A., Martinez, C., Kasavajhala, K., Wickstrom, L., Hauser, K.E., Simmerling, C., 2015. ff14SB: Improving the Accuracy of Protein Side Chain and Backbone Parameters from ff99SB. J. Chem. Theory Comput. 11, 3696–3713. https://doi.org/10.1021/acs.jctc.5b00255

Pettersen, E.F., Goddard, T.D., Huang, C.C., Meng, E.C., Couch, G.S., Croll, T.I., Morris, J.H., Ferrin, T.E., 2021. UCSF ChimeraX: Structure visualization for researchers, educators, and developers. Protein Sci. Publ. Protein Soc. 30, 70–82. https://doi.org/10.1002/pro.3943

Proceedings of the Python in Science Conference (SciPy): Exploring Network Structure, Dynamics, and Function using NetworkX [WWW Document], n.d. URL http://conference.scipy.org.s3-website-us-east-1.amazonaws.com/proceedings/scipy2008/paper_2/ (accessed 3.2.25).

Roe, D.R., Cheatham, T.E., 2013. PTRAJ and CPPTRAJ: Software for Processing and Analysis of Molecular Dynamics Trajectory Data. J. Chem. Theory Comput. 9, 3084–3095. https://doi.org/10.1021/ct400341p

Wang, J., Wang, W., Kollman, P.A., Case, D.A., 2006. Automatic atom type and bond type perception in molecular mechanical calculations. J. Mol. Graph. Model. 25, 247–260. https://doi.org/10.1016/j.jmgm.2005.12.005

Wang, J., Wolf, R.M., Caldwell, J.W., Kollman, P.A., Case, D.A., 2004. Development and testing of a general amber force field. J. Comput. Chem. 25, 1157–1174. https://doi.org/10.1002/jcc.20035
