## Supplementary material for "Musicality in protein interaction dynamics informs the multi-scale evolution of prosocial behavior": examples in .ppt

### Slide 1
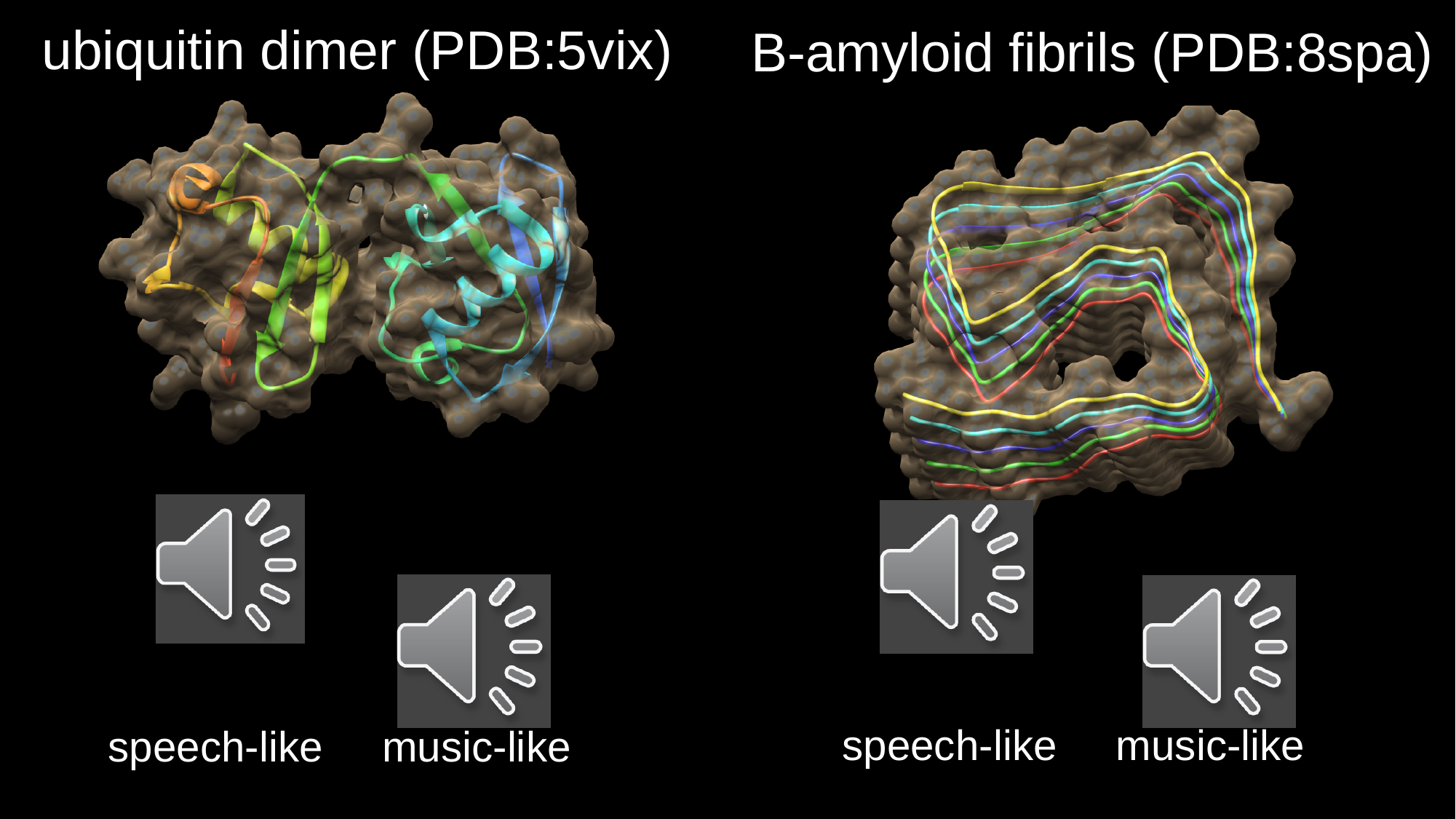

ubiquitin dimer (PDB:5vix)
B-amyloid fibrils (PDB:8spa)
speech-like music-like
speech-like music-like

### Slide 2
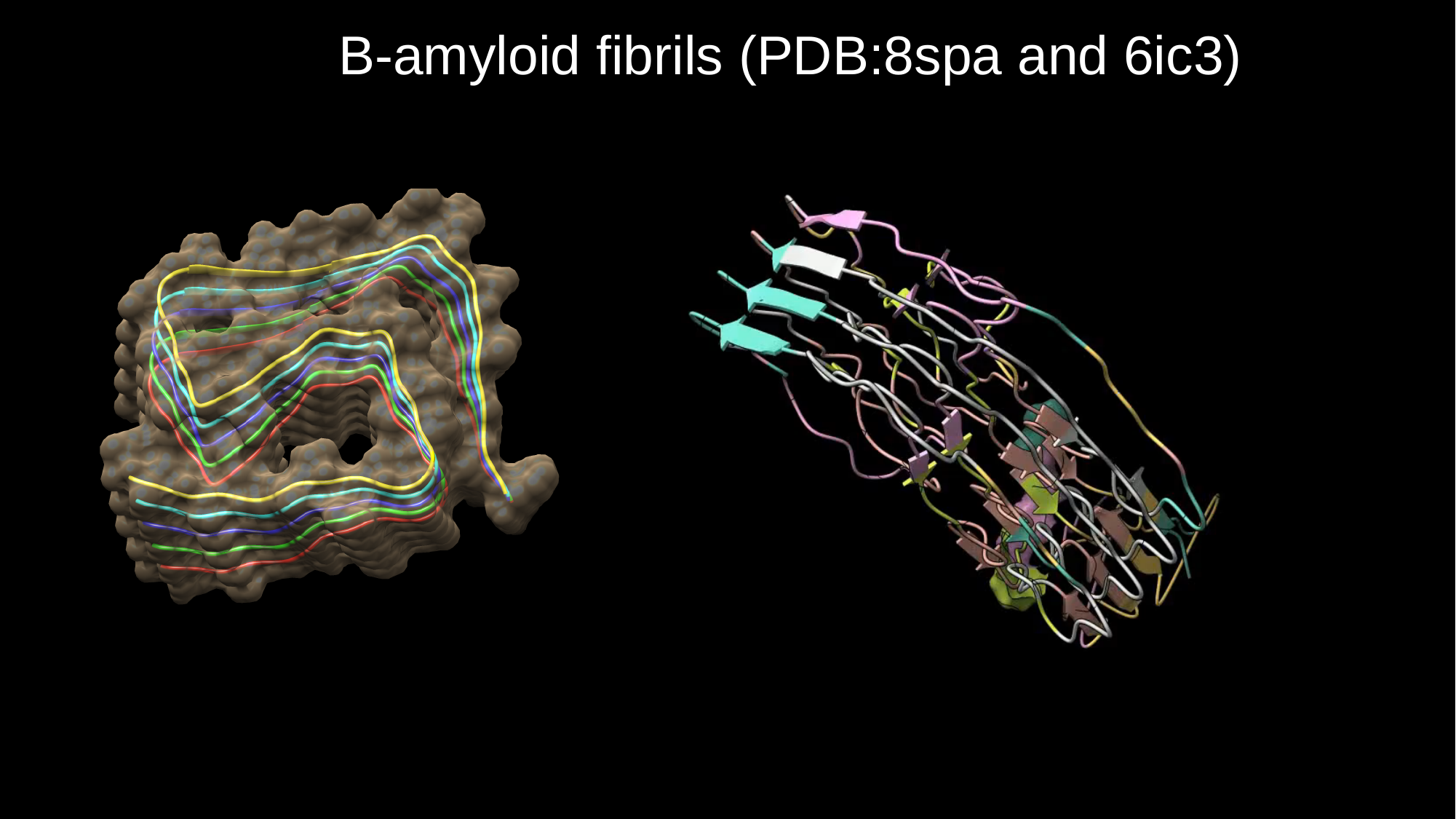

B-amyloid fibrils (PDB:8spa and 6ic3)
